# SIRT6 Activation Improves Intervertebral Disc Health in the Aging Spine

**DOI:** 10.64898/2026.03.03.709336

**Authors:** Pranay Ramteke, Bahiyah Watson, Sanjana Jagannath, Sarah E. Bell, Mallory Toci, Olivia Ottone, Qinglin Wu, Oliver Abinader, Ramkrishna Mitra, John A. Collins, Ruteja A. Barve, Makarand V. Risbud

## Abstract

Aging is one of the most important risk factors for Intervertebral disc degeneration, a major contributor to chronic low back and neck pain. Recently, we demonstrated a critical role for SIRT6, a nuclear NAD⁺- dependent deacetylase and defatty acylase, in maintaining intervertebral disc health with aging. We therefore investigated whether pharmacological activation of SIRT6 improves disc health by examining the spinal phenotype of 24-month-old mice treated with the well-studied agonist MDL-800 for 6 months. Histological studies revealed healthy disc tissue morphology, enhanced cell viability, and lower degeneration scores in mice treated with MDL-800. Further mechanistic insights revealed that SIRT6 activation decreased H3K9ac levels, improved cell phenotype and matrix quality, and reduced the SASP burden in the disc, characterized by decreased abundance of p21, IL-6, and TGF-β. Tissue RNA-Seq, *in vitro* measurements of histone 3 modifications, and multi-omics ATAC-seq/RNA-seq analyses revealed that SIRT6 activation altered the epigenetic status (decreased H3K9ac, H3K36me3, and H3K79me2) and transcriptomic landscape of disc cells. Notably, MDL-800 treatment increased LC3II levels in disc cells, indicating enhanced autophagic flux. Furthermore, plasma LC-MS and nuclear magnetic resonance (NMR) analyses revealed minimal systemic metabolomic changes. ScRNA-sequencing of splenocytes and bone marrow cells and systemic cytokine profiling indicated good tolerance and the absence of systemic inflammation following MDL-800 treatment. Our study demonstrates that SIRT6 activation modulates autophagy, cell senescence, and matrix homeostasis in the disc, underscoring the feasibility of targeting SIRT6 activation as a promising pharmacological strategy to maintain disc health in the aging spine.

## Introduction

Low back pain (LBP) is the leading cause of global disability, affecting an estimated 619 million individuals in 2020, with projections reaching 843 million by 2050 due to population growth and aging (*1*). The peak prevalence of LBP is observed between the ages of 50 and 55, extending into older age groups (*1*). One of the primary contributors to LBP is intervertebral disc degeneration and age-related spinal changes, which lead to structural degeneration, pain, and functional impairments. Disc degeneration is a progressive, multifactorial process involving dysregulated autophagy, cellular senescence, chronic inflammation, and extracellular matrix degradation (*2*). Current treatments remain symptomatic and often ineffective, underscoring the need for non-invasive, biologically targeted disease-modifying strategies to reduce the growing global burden of LBP.

Sirtuin 6 (SIRT6) is a nuclear NAD⁺-dependent enzyme with deacetylase, defatty-acylase, and mono-ADP-ribosyltransferase activities. By removing acetyl groups from histone marks such as H3K9ac, H3K18ac and H3K56ac, SIRT6 preserves chromatin integrity and coordinates DNA repair, metabolic regulation, and inflammatory responses - processes essential for tissue homeostasis and healthy aging (*3–6*). Recent findings from human lifespan studies show a prominent role of SIRT6 in aging and age-associated disorders (*7, 8*). In mouse models, loss of SIRT6 accelerates genomic instability, metabolic dysfunction, premature aging, and reduced lifespan, disproportionately affecting males (*9, 10*). While Kanfi et al. reported a significantly longer lifespan in male SIRT6 transgenic mice through downregulation of IGF-1 signaling, a later study showed lifespan extension in both male and female mice, albeit with a stronger effect in males than in females via preservation of hepatic glucose output and glucose homeostasis (*11, 12*). Similarly, a positive impact of SIRT6 on lifespan in other species, involving diverse mechanisms such as efficient DNA double-strand break repair and MYC inhibition, has been demonstrated, underscoring SIRT6’s role as an epigenetic regulator of genome stability and aging (*13, 14*).

SIRT6 plays a prominent role in the musculoskeletal system (*15, 16*). *Sirt6*^AcanCreERT2^ mice (Sirt6 loss) show significantly repressed IGF-1/Akt signaling that was linked to enhanced injury-induced and age-associated osteoarthritis (OA) severity (*17, 18*). Moreover, these SIRT6-deficient mouse chondrocytes showed diminished antioxidant capacity, evidenced by reduced peroxiredoxin 1 (Prx1) and increased thioredoxin interacting protein (TXNIP) levels (*18*). Similarly, in another mouse model of SIRT6-loss in cartilage (*Sirt6*^Col2a1CreERT2^) increased chondrocyte senescence and age-associated OA severity, and a critical role of SIRT6 in STAT5 deacetylation, inhibiting the pathogenic IL-15/JAK3/STAT5 axis was noted (*19*). Notably, SIRT6 activation in chondrocytes suppresses age-related DNA damage (*20*). Loss of SIRT6 in osteoblast lineage cells resulted in reduced osteoprotegrin levels and increased osteoclast activation, leading to osteopenia (*15*). Likewise, osteoblast- and osteocyte-targeted *Sirt6*^Dmp1Cre^ mice exhibited osteopenia due to elevated osteocytic expression of *Sost* and *Fgf23,* as well as increased cell senescence (*21*). An inactivating SIRT6 mutation has been linked to perinatal lethality and craniofacial abnormalities in humans (*22*), and *Sirt6* deletion in Wnt1-expressing neural crest cells in mice delays palatal osteogenesis, resulting in a cleft palate (*23*). In hematopoietic cells, SIRT6 promotes the osteoclast differentiation program by increasing NFATc1 and Blimp1, and by suppressing MafB (*24*). In the rotator cuff, an age-related decline in SIRT6 has been postulated to contribute to impaired tendon-to-bone healing after injury. Experimental overexpression in rotator cuff and primary tenocytes restored Wnt/β-catenin signaling and reduced sclerostin expression, improving enthesis repair in aged rats following tenotomy with reconstruction, suggesting therapeutic potential for tendon injuries in older individuals (*25*). These context-specific roles of SIRT6 underscore its importance in maintaining skeletal tissue homeostasis.

In our recent studies, disc-targeted Sirt6 deletion in mice disrupted disc homeostasis and increased degenerative features as the animals aged. We observed increased H3K9ac, DNA damage, cell senescence, and the senescence-associated secretory phenotype (SASP), accompanied by decreased cellular autophagy, in knockout mice, indicating a causal link between SIRT6 activity and disc health (16). Similarly, SIRT6 activation has been shown to suppress senescence in an acute traumatic disc injury model (26). Whether it will improve disc health in the context of age-dependent degeneration remains unknown. Preclinically validated SIRT6 activators have been shown to accelerate wound healing (27), mitigate hepatic fibrosis (28, 29), and improve muscle performance (30), demonstrating their utility for in vivo applications. We treated aged B6 mice with MDL-800, a well-studied SIRT6 allosteric activator, and observed a significant improvement in disc health outcomes compared to vehicle-treated, age-matched mice. Mechanistically, there were changes in chromatin accessibility, evident by reduced H3K9 acetylation, decreased cell death, senescence, and increased autophagy. Systemically, we observed minimal alterations in the plasma cytokine levels and the metabolome, accompanied by few changes in the immune cell profile, suggesting an anti-inflammatory phenotype. These findings suggest that pharmacological activation of SIRT6 by MDL-800 may represent a promising biological therapeutic strategy for treating age-related disc degeneration.

## Results

### MDL-800 treatment improves disc health in old mice without adverse effects on systemic metabolism

To investigate whether SIRT6 activation improves disc health in the context of age-dependent degeneration, we examined the spinal phenotype of B6 mice treated with MDL-800 for 6 months starting at 18 months of age (Fig. 1A). MDL-800 treatment significantly decreased H3K9 acetylation levels in the disc tissues, confirming increased SIRT6 activity (Fig.1B). When disc degeneration was assessed histologically in the lumbar spine, treatment with MDL-800 lead to a marked decline in the age-dependent degenerative features, including NP fibrosis, loss of NP-AF compartment demarcation, lamellar defects, and clefts indicative of structural disruptions (Fig. 1C). Notably, NP cells showed a healthy morphology as evident from preservation of vacuolated cells in the cell band. Modified Thompson grading of MDL-800 treated mice showed significantly lower scores of degeneration in both NP and AF compartments compared to vehicle-treated controls (Fig. 1C-E). Since cell death is observed during disc degeneration, we performed TUNEL labelling, which showed a significant decrease in TUNEL-positive cells in MDL-800 treated mice, but a comparable total cell number, indicating that the elevated apoptosis was not the primary driver of later stages of degeneration (Fig. 1F,G). Despite an overall improvement in disc health parameters, MDL-800 treated mice did not exhibit altered disc height (DH) or disc height index (DHI) when compared to vehicle-treated controls (Fig. 1H-K).

**Figure 1.**
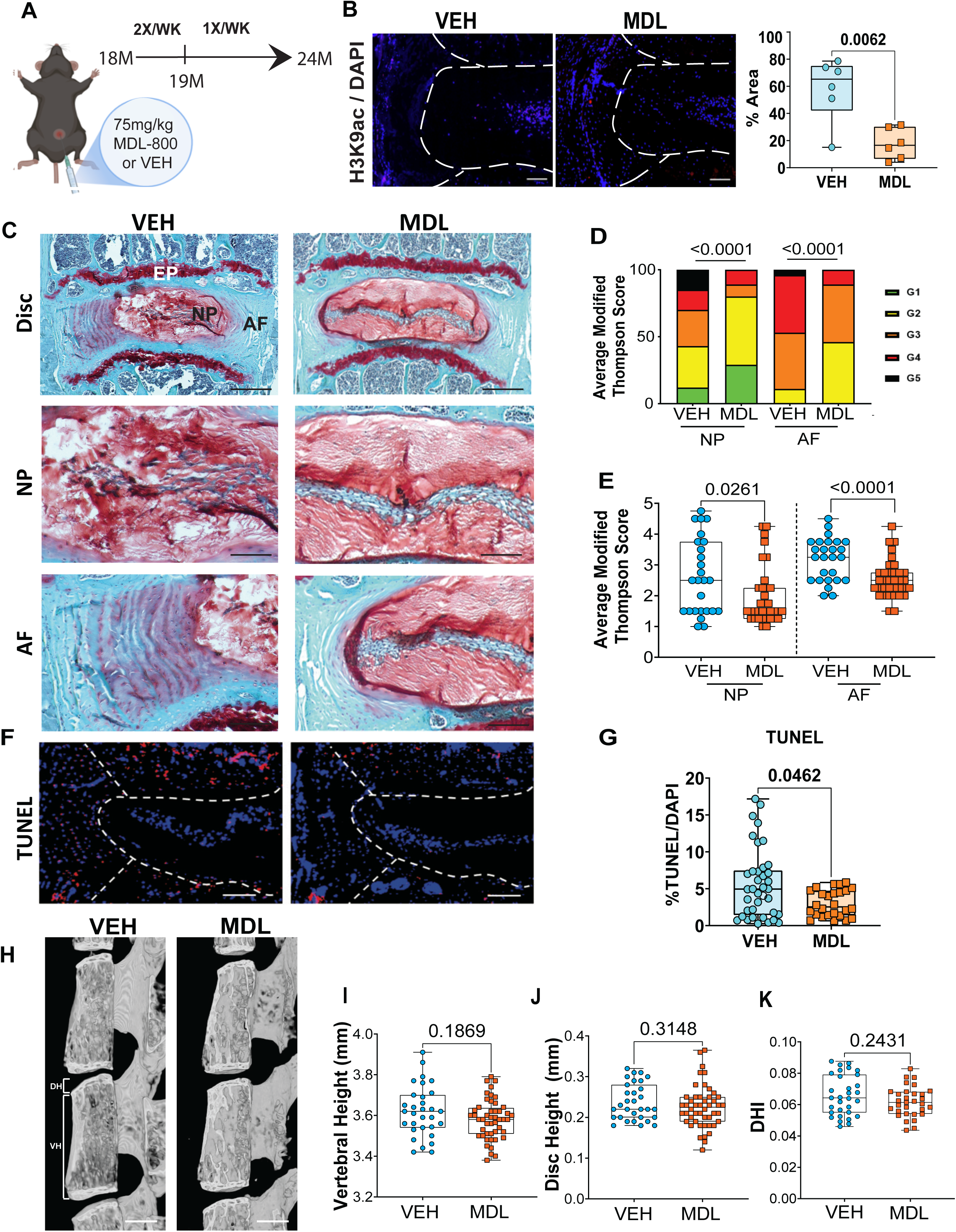
SIRT6 activation by MDL-800 improves age-associated intervertebral disc health. (A) Experimental design showing the timeline of MDL-800 treatment of 18-month-old B6 mice. (B) Immunofluorescence staining for H3K9ac shows a significant decrease in staining in the lumbar disc, confirming SIRT6 activation. (C) Safranin-O/Fast Green staining of lumbar discs of vehicle control (VEH) and MDL-800 (MDL) treated mice at 24 months. Scale bar row 1 = 200 μm; row 2 = 50 μm. (D, E) Distribution of and average modified Thompson’s grades for NP and AF compartments of lumbar discs of VEH and MDL-800 mice, n = 7 Vehicle Control, and 9 MDL-800 mice, 2-4 discs/animal. (F) TUNEL staining images and (G)TUNEL quantification, Scale bar = 50 μm. (H-K) μCT analysis showing disc height (DH), vertebral height (VB), and disc height index (DHI) measured at 24 months. n = 8-12 mice/group, 3 discs/animal, scale bar = 1 mm. Quantitative measurements are reported as the median with the interquartile range. Statistical difference between grade distributions (D) was tested using the chi-square test, and all other quantitative data were compared using an unpaired t-test or Mann–Whitney test, as appropriate.

We then performed plasma metabolic analyses to determine the systemic effects of long-term MDL-800 treatment in old mice using LC/MS (Fig.S1) and NMR (Figs. S2 and S3) assays. LC/MS analysis identified 184 plasma metabolites with high confidence but did not reveal any treatment-dependent PCA or hierarchical clustering (Fig. S1B, C). Out of 6 metabolites that showed significant differences, 4 metabolites, including palmitoyl sphingomyelin, 5-aminolevulinic acid, 2-oxindole, and jasmonic acid, showed a significant increase, while nicotinamide and hypotaurine showed a decrease in MDL-treated mice (Fig. S1D-I). Plasma NMR analysis showed no changes in any of the measured analytes, including indicators of inflammation and metabolic regulation: GlycA, Total Ketone Bodies, Acetoacetate, Acetone, Beta Hydroxy-butyrate Total branched chained amino acids (BCAA), Alanine, Leucine, Isoleucine, Valine, Citrate, ApoA-1, ApoB, Total Triglyceride, Total Cholesterol, Total TRLP, Small TRLP, total calibrated low-density lipoprotein particle (cLDLP), total calibrated high-density lipoprotein particle (cHDLP), medium LDLP, H4P, TRL Size, LDLC, HDLC, HDL Size, TRL TG (Fig. S2B-M and S3). In conclusion, these plasma analyses revealed minimal metabolic effects of long-term MDL-800 treatment in B6 mice, supporting a favorable safety profile under the conditions tested.

### MDL-800 treatment preserves NP cell phenotype, aggrecan levels, and ameliorates disc fibrosis in aged mice

As SIRT6 activation improved histological scores of disc health, with a notable effect on NP cell morphology, we determined whether MDL-800 treatment preserved cell phenotype and matrix constituents using Picrosirius red staining and quantitative immunohistochemistry. Polarized imaging of picrosirius red-stained disc sections showed diminished presence of collagen fibers in the NP compartment and an increased abundance of small-diameter fibers and a decrease in thick fibers in the AF of MDL-800 treated mice (Fig.2A-D). These findings suggest an overall increase in collagen turnover and amelioration of age-associated NP fibrosis resulting from sustained SIRT6 activation. To determine the integrity of the disc ECM, we stained the disc sections for Aggrecan (ACAN), a predominant constituent of the NP matrix, and noted a significant increase in abundance (Fig.2E, F). Likewise, NP cells showed increased GLUT1 abundance, a known phenotypic marker, indicating preservation of NP tissue health (Fig. 2G, H). There were little changes in the abundance of COL1 and FCHP, a marker of denatured collagen in the AF (Fig. S4). Similarly, both groups exhibited comparable COLX abundance in the NP (Fig. S4), suggesting that SIRT6 activation did not modulate the age-dependent acquisition of chondrocyte-like characteristics. These findings suggested that SIRT6 is critical in maintaining a healthy cell phenotype and disc tissue matrix.

**Figure 2.**
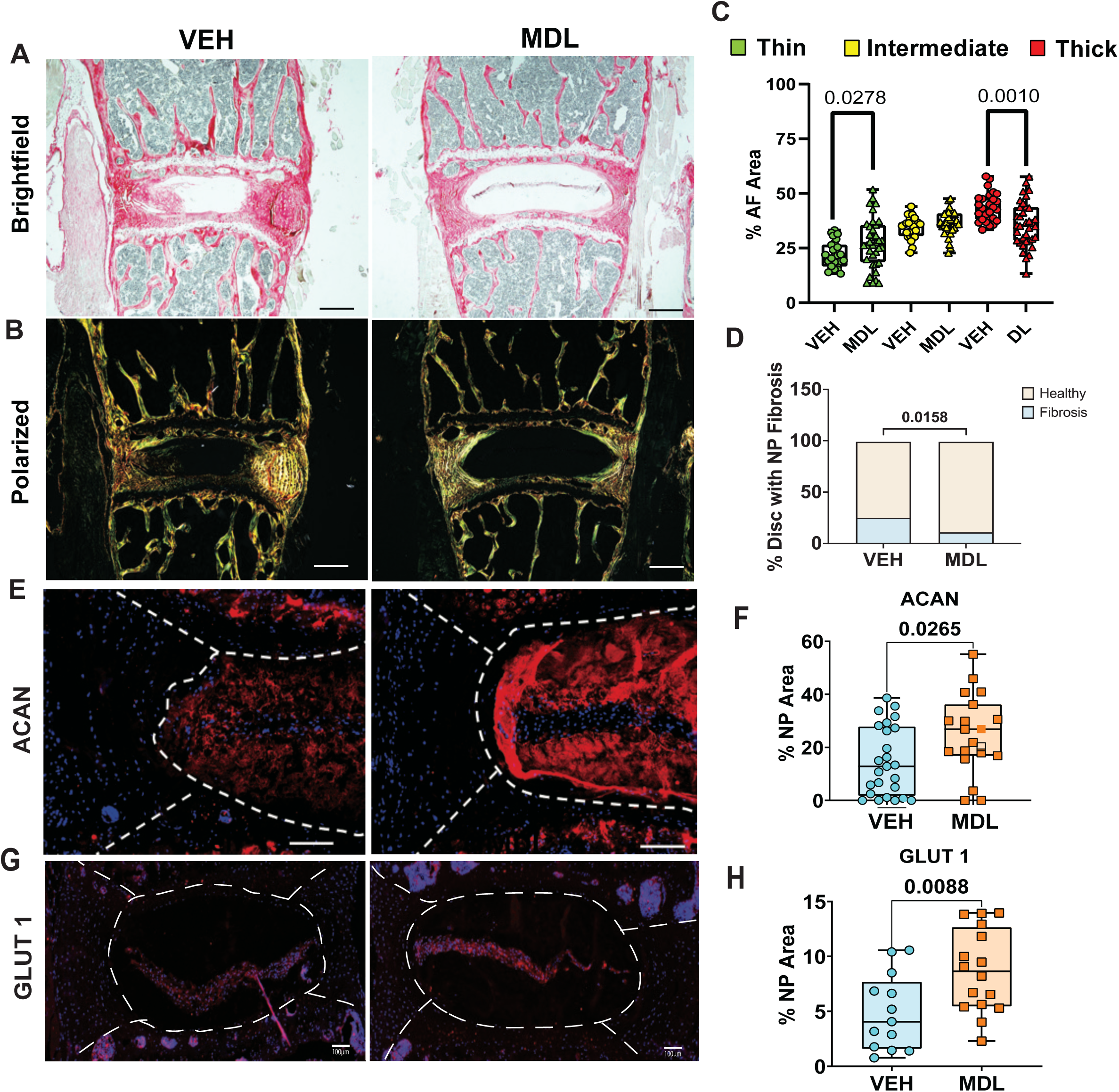
SIRT6 activation promotes ECM homeostasis and decreases NP tissue fibrosis. (A) Representative bright field and (B) polarized images of Picrosirius Red-stained lumbar disc sections and (C) quantification of collagen fibers from VEH and MDL-800 treated mouse discs, n = 7 Vehicle Control, and 9 MDL-800 mice; 2-4 discs/mouse. Scale bar= 200 μm. (D) Quantification of NP compartment fibrosis using polarized images. (E-F) Representative immunofluorescence images of lumbar disc sections and (G, H) corresponding quantification for Aggrecan (ACAN) and GLUT1. Scale bar = 50 μm, n = 6 mice/group and 2-4 discs/animal. White dotted lines demarcate disc compartments. Quantitative measurements are reported as the median with the interquartile range, and significance was determined by using an unpaired *t*-test or Mann–Whitney test, as appropriate.

### SIRT6 activation diminishes senescence and SASP burden in the disc

The senescence burden and SASP have been causally linked to age-dependent disc degeneration (*31*). We therefore investigated the senescence status of MDL-800 treated mice. We noted that levels of p21, a Cdk inhibitor important for senescence initiation, were significantly decreased in the discs of MDL-800 treated mice (Fig.3A, B) (*32*). Additionally, we determined levels of IL-6 (Fig.3C, D) and TGF-β (Fig.3E, F), important markers of SASP in the disc (*31*). Notably, IL-6 and TGF-β levels were significantly lower in discs from MDL-800 treated mice, suggesting a lower overall burden of SASP and profibrotic factors.

**Figure 3.**
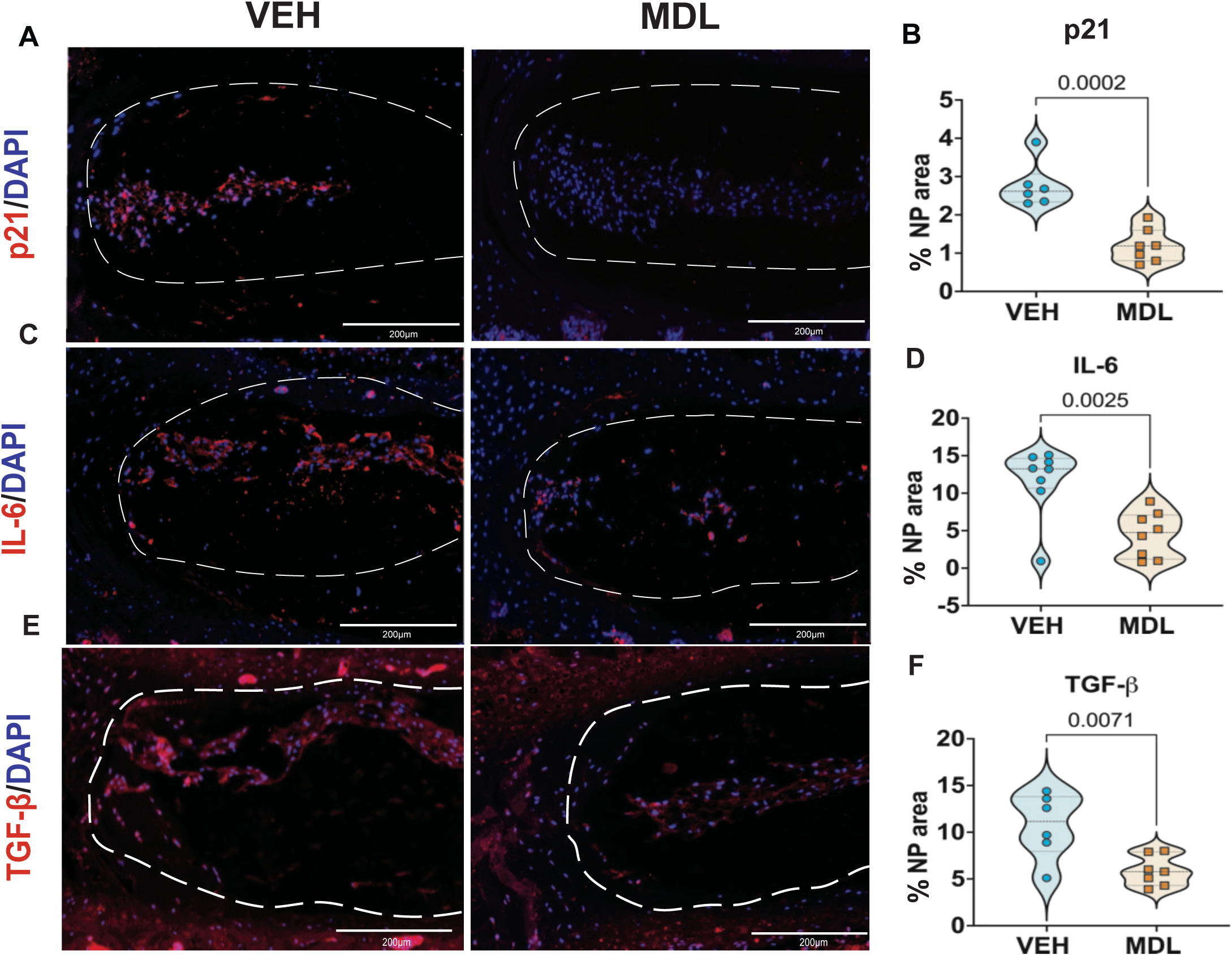
SIRT6 activation decreases senescence and SASP burden in intervertebral discs of MDL-treated mice. (A-F) Representative immunofluorescence images and corresponding quantitative analysis of senescence and SASP markers (A, B) p21, (C, D) IL-6 and (E, F) TGF-β, n = 4 animals/group, 3 discs/mouse. Scale bar = 50 μm.

### SIRT6 activation drives changes in the transcriptomic profile of NP and AF tissues

To determine the broader effects of long-term, sustained SIRT6 activation on the transcriptomic landscape in NP and AF tissues, we performed RNA sequencing. While SIRT6 suppresses gene expression via limiting chromatin accessibility, long-term effects were expected to reflect second-order adaptive changes. Indeed, we observed that more genes were upregulated than downregulated in both NP and AF tissues by MDL-800 treatment (Fig. S5). In NP tissue, there were major upregulated thematic clusters related to 1) Histone acetylation/activity, and JNK/MAPK signaling 2) a large supercluster for circadian clock, cell polarity/notochord, follistatin-inhibin-activin signaling, connexins/Cx43, and matrix remodeling 3) lipid metabolism and PPAR signaling, and 4) microtubules anchoring at centrosome (Fig. 4). Notably, when downregulated DEGs were analyzed, we observed a less prominent thematic clustering with major clustering centered around 1) follistatin-myostatin and actin-myosin, 2) connexins/Cx43 and Natriuretic peptide/myocardium (Fig. S6). The upregulated DEGs in AF clustered into two major superclusters. The first cluster was related to skeletal tissues, with themes including articular cartilage development, endochondral processes, and collagen/MMPs. While the second supercluster was related to neuronal processes, it encompassed themes such as the hypothalamic nucleus, synaptic vesicle exocytosis, axonal guidance, sensory neurons, synapses, clathrin-coated vesicles, and voltage-gated Ca^2+^ channels (Fig. S7). Again, compared with the extensive clustering observed in upregulated DEGs in AF, less clustering was observed in downregulated DEGs. Some of the enriched themes in these downregulated DEGs included immune cell function (CD8 effector T cells, granzyme secretion, MCP-1 Production, PD-1 secretion, macrophage response), caspase activation, myogenesis, Schwann cells, and oligodendrocytes and myelination (Fig. S8).

**Figure 4.**
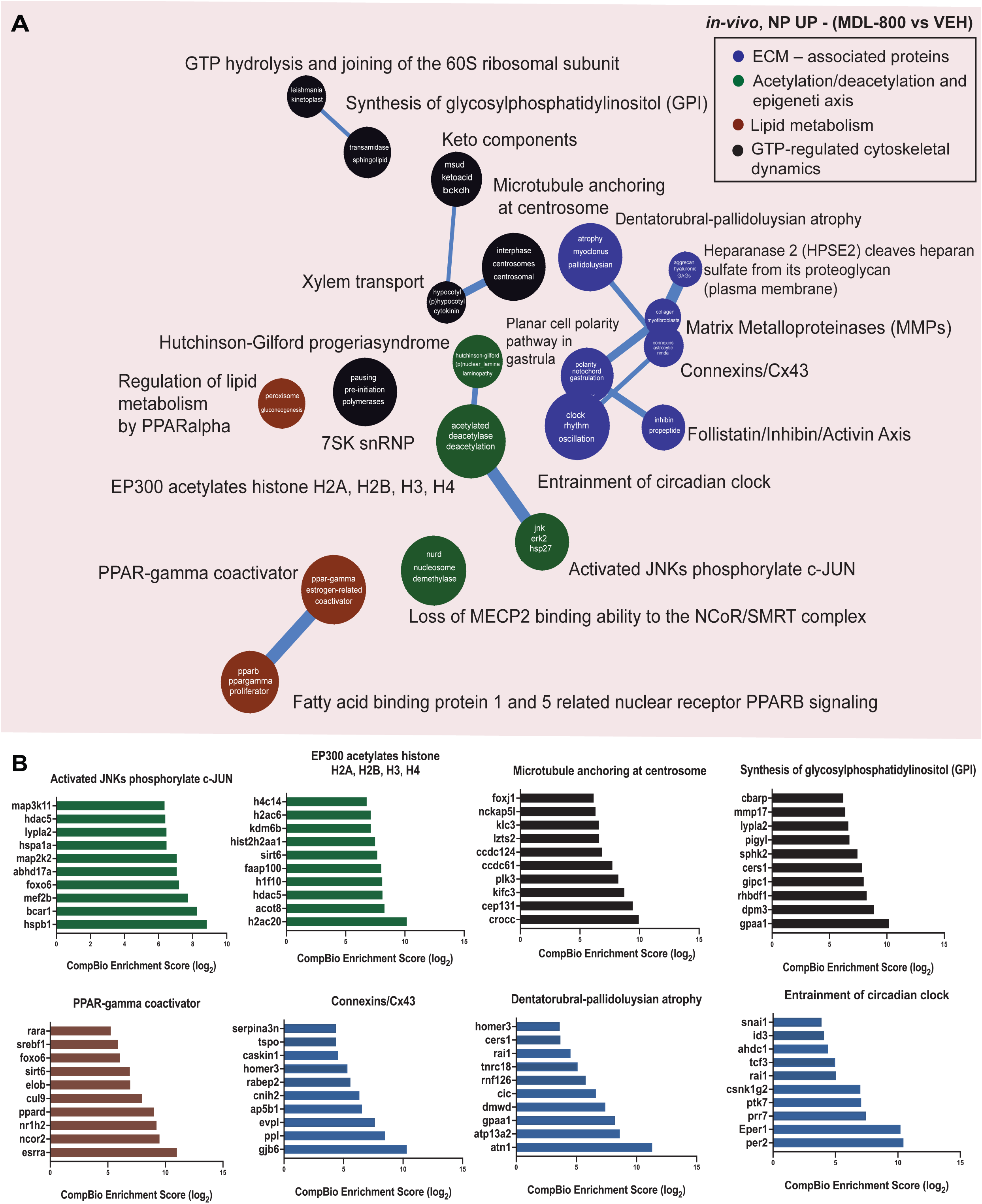
SIRT6 activation alters the transcriptomic landscape of NP tissue. (A) Thematic map of CompBio analysis of Upregulated DEGs (p < 0.05, FC > 1.75) from RNA sequencing of NP tissue transcripts from 24-month-old, MDL-800 treated B6 mice compared to VEH-treated mice, represented as a ball and stick model, n = 5 mice/group, 5-6 pooled discs/animal. DEGs were mapped to major themes related to Histone acetylation, JNK/MAPK signaling, circadian clock, follistatin/inhibin, connexins, matrix remodeling, lipid metabolism, and microtubules. The enrichment of themes is shown by the size of the ball, and the connectedness between themes is proportional to the thickness of the lines between them. Themes of interest are colored, and superclusters comprised of related themes are highlighted. (B) Top thematic DEGs plotted based on CompBio entity enrichment score.

### MDL-800 treatment causes changes in immune cell profiles in lymphatic tissues without systemic inflammation

Immune cells play a crucial role in regulating aging phenotypes (*33*). To investigate the immune cell profile and its association with disc phenotype in MDL-800 treated mice, we performed single-cell RNA sequencing (scRNA-seq) of two major lymphatic tissues, the spleen and bone marrow (Fig.5, S9, S10). After normalization, the splenocytes (Fig. 5) and bone marrow cells (Fig. S9) were grouped into 12 and 15 clusters, respectively, using the FindNeighbors and FindClusters functions in the Seurat single-cell analysis package. Moreover, the data was visualized according to treatment, and cell populations were identified using the SingleR annotation tool, demonstrating differences in the proportion of B cells, T cells and granulocytes and a small change in monocytes or macrophages in the MDL-800 treated group in lymphoid tissues (Fig. 5B-E, and Fig. S9B-E). Previous studies have implied a relationship between immune cell heterogeneity with functional state and origin (*34, 35*). To further visualize cellular heterogeneity, T cell populations were subdivided into CD4+, CD8a+, and CD4-, CD8a- T cells (Fig. S10). We observed an overall decrease in T cell numbers in the MDL-800-treated mice (Fig. S10A-D, G-J). When cell proportions were compared, there was a relative increase in the proportion of CD4+ and a decreased proportion of CD8a+, and CD4-, CD8a- T cells in the spleen (Fig. S10D’). However, there was no appreciable change in the proportion of T cell subpopulations in bone marrow (Fig. S10J’).

**Figure 5.**
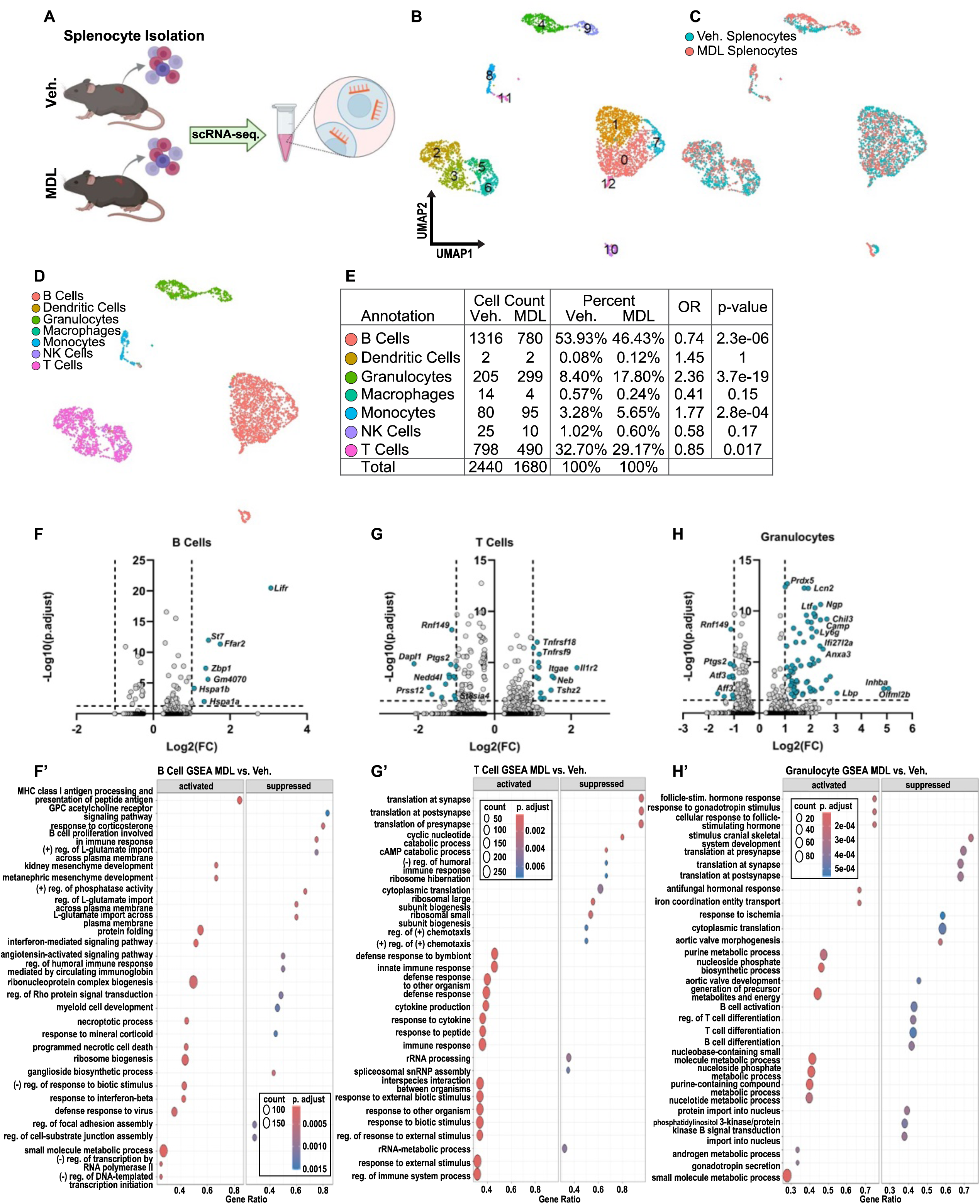
MDL-treated mice exhibit changes in splenic immune cell profiles. (A) Schematic showing the isolation and scRNA-sequencing of splenocytes from Vehicle- and MDL-800 treated mice. (B) UMAP showing unbiased clustering of splenocytes using Seurat. (C) UMAP with the annotation of Seurat clusters according to treatment. (D) UMAP with the annotation of cells using SingleR. (E) Number and proportion of cells identified in each treatment cohort, with odds ratios and p-values. Volcano plots showing differential gene expression and GSEA in (F, F’) B Cells, (G, G’) T Cells, and (H, H’) Granulocytes based on the comparison of MDL-800 to vehicle-treated splenocytes.

Gene set enrichment analysis (GSEA) was then performed on DEGs ( *P* < 0.05) for cell populations that exhibited significant differences in proportion using the clusterProfiler package (*36*). In the MDL-800 group, DEGs in splenic B cells showed activation pathways related to MHC class I processing, plasma membrane protein folding, ribonucleoprotein complex and ribosome biogenesis, and small molecule metabolic process (Fig. 5F, F′). In B cells, GSEA identified suppression of the response to corticosterone, B cell proliferation in immune response, plasma membrane L-glutamate import, and regulation of humoral immune response, with top DEGs including *Lifr*, *St7*, *Ffar2*, *Hspa1a/1b,* responsible for this enrichment (Fig. 5F, F′). In the MDL-800 group, DEGs from splenic T cells showed activation of defense response–related pathways, with top DEGs including *Tnfrsf9/18*, *Itgae*, *Il1r2,* and *Tshz2* (Fig. 5G, G’). T cells also exhibited DEGs linked to the suppression of multiple pathways related to protein translation, ribosomes, and rRNA, with top DEGs that included *Dapl1*, *Nedd4l, Prss12*, and *Ptgs2* (Fig. 5G, G’). GSEA of granulocytes showed that MDL-800 treatment led to activation of the reproductive hormone response and nucleoside metabolism, suppression of pathways related to translation, ischemic response, and an overall decrease in B- and T-cell activation and differentiation (Fig. 5H, H’). There was no appreciable enrichment of DEGs in GSEA pathways in bone marrow immune cells in MDL-800 treated mice (Fig. S9).

To investigate whether MDL-800 dependent changes in immune cell profiles in primary lymphatic tissues were correlated with systemic inflammation status, we measured plasma levels of key cytokines and chemokines. We did not observe any significant alterations in levels of IFNγ, IL-1β, IL-2, IL-4, IL-5, IL-6, IL-10, IL-12/p70, IL-15, IL-17A/F, IL-27/p28/IL-30, IL-33, IP-10, KC/GRO, MCP-1, MIP-1α, MIP-2, and TNF-α (Fig.S11A-R), suggesting an absence of systemic inflammation.

### SIRT6 activation alters H3K modifications and increases autophagy in NP cells without affecting cellular metabolism

To delineate mechanistic insights into how MDL-800 treatment and subsequent SIRT6 activation affect disc cells, we performed *in vitro* experiments using primary rat NP cells (Fig.6A). Western blot analysis showed a significant decrease in H3K9ac, a known marker of SIRT6 activation, without changes in SIRT6 levels in MDL-800 treated NP cells (Fig.6B, C). This finding is further supported by immunofluorescence staining showing decreased nuclear staining of H3K9ac and comparable levels of H3K27ac in MDL-800-treated cells (Fig. 6D).

**Figure 6.**
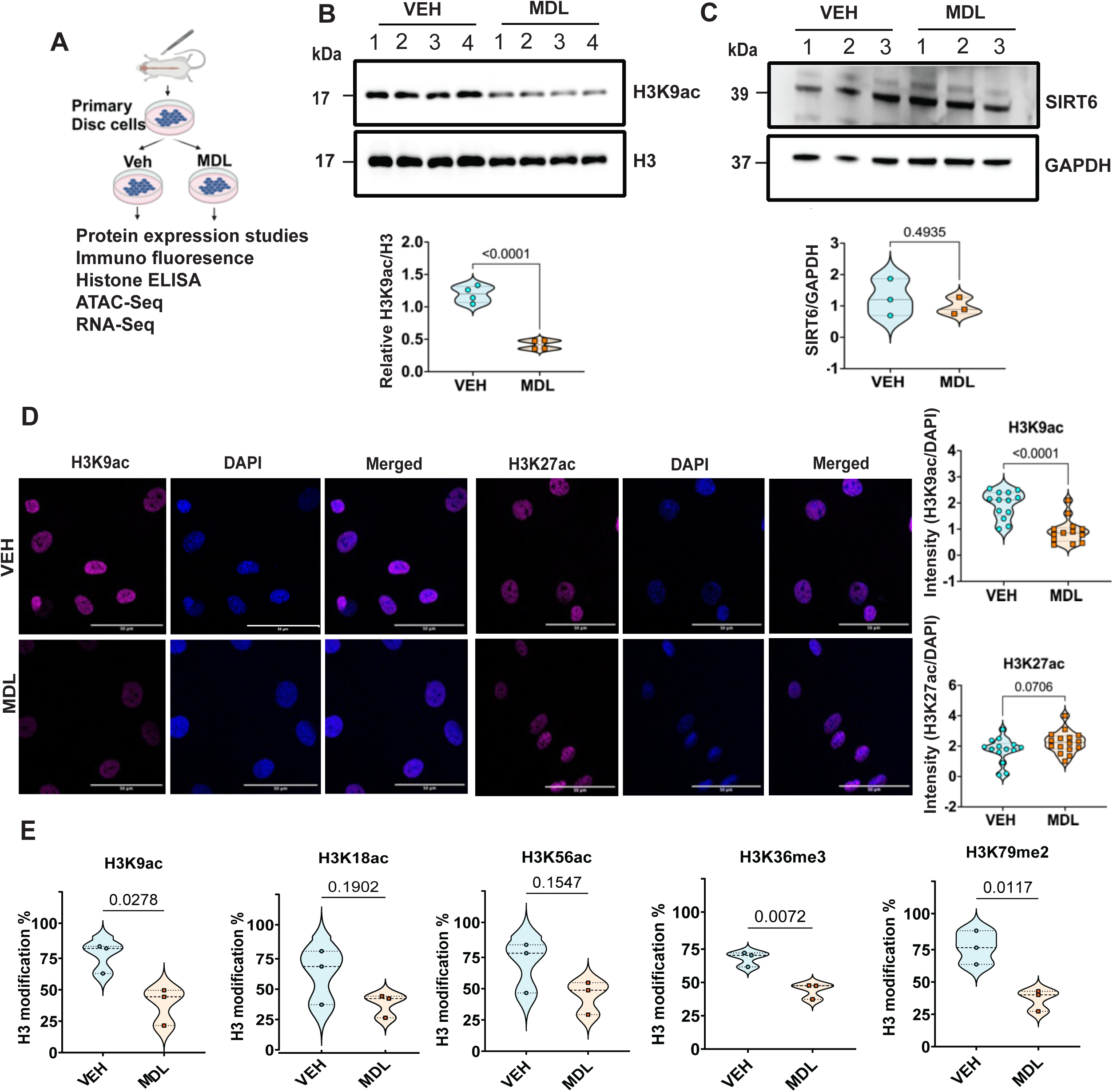
SIRT6 activation alters histone modifications and chromatin accessibility in disc cells. (A) Schematic of *in vitro* experimental design. (B, C) Western blot analysis of (B) H3K9ac and (C) SIRT6 and corresponding densitometric quantitation normalized to total H3 and GAPDH of VEH and MDL-800 treated NP cells. n = 4 independent cell isolations. (D) Immunofluorescent staining and quantification of H3K9ac and H3K27ac in VEH and MDL-treated NP cells. (E) Quantitative ELISA for H3 lysine modifications from VEH and MDL-800 treated NP cells. n = 3 independent cell isolations. Statistical significance was determined by an un-paired t-test

To understand the broader consequences of SIRT6 activation, we measured various H3K modifications using an ELISA assay. We again saw a significant decrease in H3K9 acetylation levels in MDL-treated NP cells (Fig. 6E). Moreover, in line with our earlier studies, acetylation status of H3K18 and H3K56 was unaffected (Fig. 6E). MDL-800 treatment also decreased methylation of H3K36me3 and H3K79me3 in NP cells without changes in H3K4me, H3K9me, H3K27me, H3K36me1, H3K36me2, H3K79me1 and H3K79me3 levels (Fig.6E and S12), underscoring tissue-type specificity in epigenetic changes following elevated SIRT6 action.

SIRT6 modulates autophagy in various tissues, including disc cells (*16, 37*), and plays a crucial role in maintaining disc health (*2*). To investigate whether SIRT6 activation induces autophagy, we measured autophagic flux in the presence of MDL-800 and Bafilomycin A1, a known autophagy inducer. Similar to Bafilomycin A1, treatment of cells with MDL-800 resulted in a significant increase in LC3II levels and the LC3II/LC3I ratio, suggesting increased autophagic flux (Fig.7A, B). Furthermore, we observed a significant increase in the number of LC3-positive autophagic puncta in MDL-800-treated NP cells (Fig.7C, D). Since SIRT6 is known to modulate glycolytic metabolism, we determined the effect of MDL treatment on the bioenergetics of NP cells using Seahorse assays (*38*). MDL-800-treated cells showed ATP production rates comparable to those of vehicle-treated control cells, suggesting no effect on cellular bioenergetics in the short term (Fig. 7E-G).

**Figure 7.**
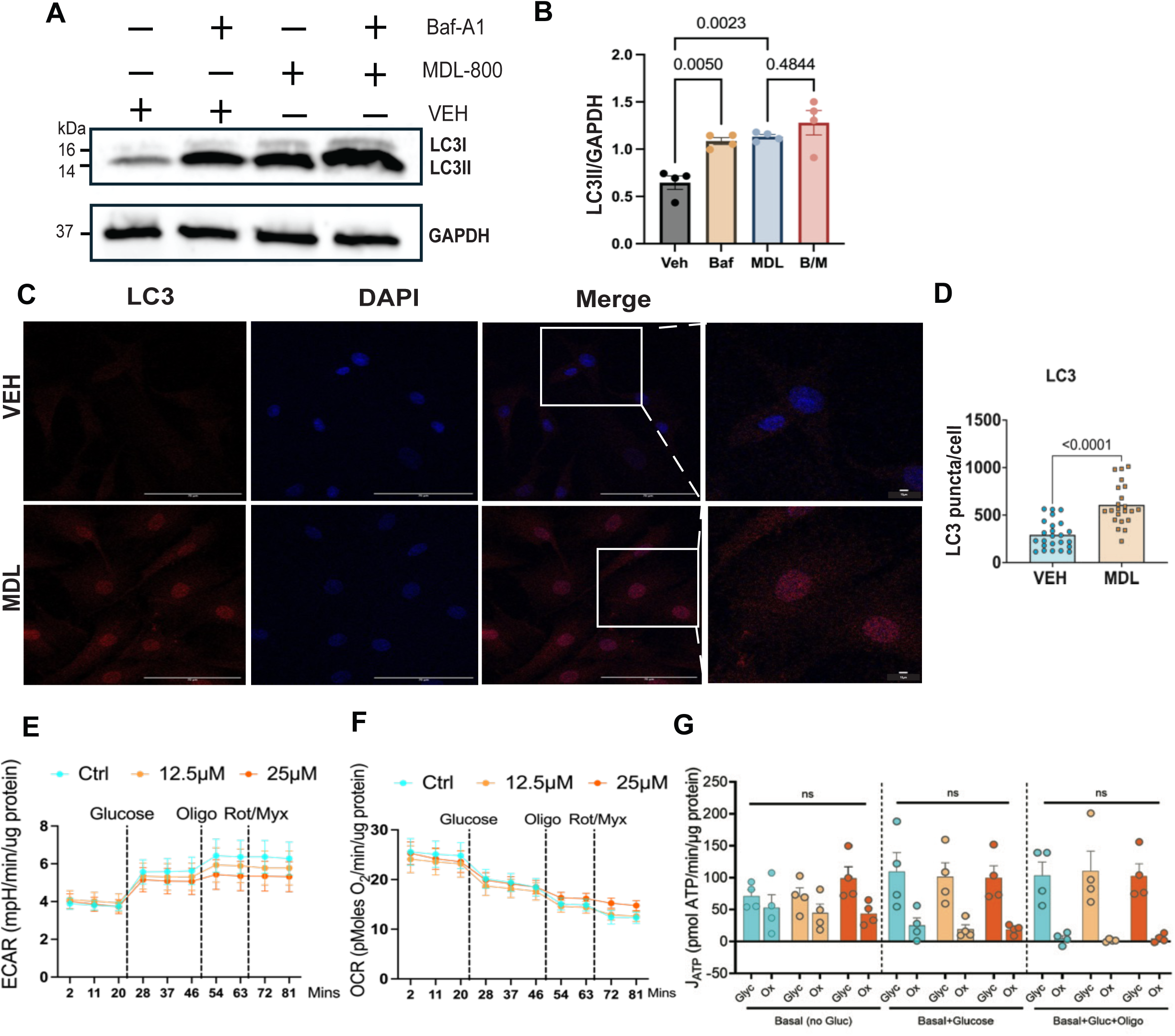
SIRT6 activation promotes NP cell autophagy without altering cellular bioenergetics. (A) Immunoblot of LC3 and (B) corresponding densitometric quantitation of LC3II/GADPH of VEH and MDL-800 treated NP cells with or without BafA1. n = 4 independent cell isolations. (C) Immunofluorescence analysis and (D) quantitation of LC3 puncta in VEH and MDL-800 treated NP cells cultured in hypoxia. Significance was tested with an unpaired t-test. Scale bar = 70 μ. (E, F) OCR and ECAR traces in VEH Vs. NP cells treated with MDL-800 for 24 hours were used to determine the (G) ATP production rate. Quantitative data are represented as the mean ± SEM; n = 4 independent cell isolations, each experiment performed in quadruplicate. Significance was determined using one-way ANOVA with Sidak’s or Kruskal-Wallis with Dunn’s multiple comparison test.

### Transcriptomic changes in MDL-treated primary NP cells align with processes affected in mice

We performed RNA-seq and ATAC-seq to understand the direct consequence of SIRT6 activation on the transcriptomic landscape of MDL-800-treated NP cells. CompBio analysis of DEGs from RNAseq (Log_2_FC ≥1.5, FDR ≤ 0.05) revealed upregulation of three major interconnected super thematic clusters related to i) actin-myosin cytoskeleton and cell junctions, ii) skeletal tissues, TGFβ/actin/BMP signaling, and extracellular matrix, iii) glycolysis, purinergic signaling, and aryl hydrocarbon receptor activity (Fig. 8). The downregulated clusters in MDL*-*treated cells included two superclusters with themes related to: 1) fatty acid β-oxidation, PPAR signaling, mevalonate pathway, dioxygenases and diacylglycerol formation, and 2) Hsp70/Hsc70 chaperons, Jagged1-Notch1 signaling, Cx36 gap junctions, and primary cilia (Fig.S13). These processes have been correlated with disc health (*39, 40*). We then used an advanced feature of the CompBio tool, which identifies significantly affected shared pathways between two datasets, to further identify the pathways commonly affected in NP tissue from MDL-800 treated mice and in primary NP cells *in vitro*, regardless of the duration of SIRT6 activation. A significant overlap in upregulated themes was noted between the two datasets for ECM proteins, collagens, ECM turnover, MMPs, myofibroblasts, gap junctions, cell-cell and cell-matrix adhesion, actin cytoskeleton, microtubules, activin/BMP/FGF, and eNOS signaling (Fig. S14A). Within downregulated datasets, themes related to ECM, collagens and fibronectin, chemokine and cytokines (interferon, IL12), coagulation and clotting (fibrinogen, fibrinolysis, and plasmin) were shared (Fig. S14B).

**Figure 8.**
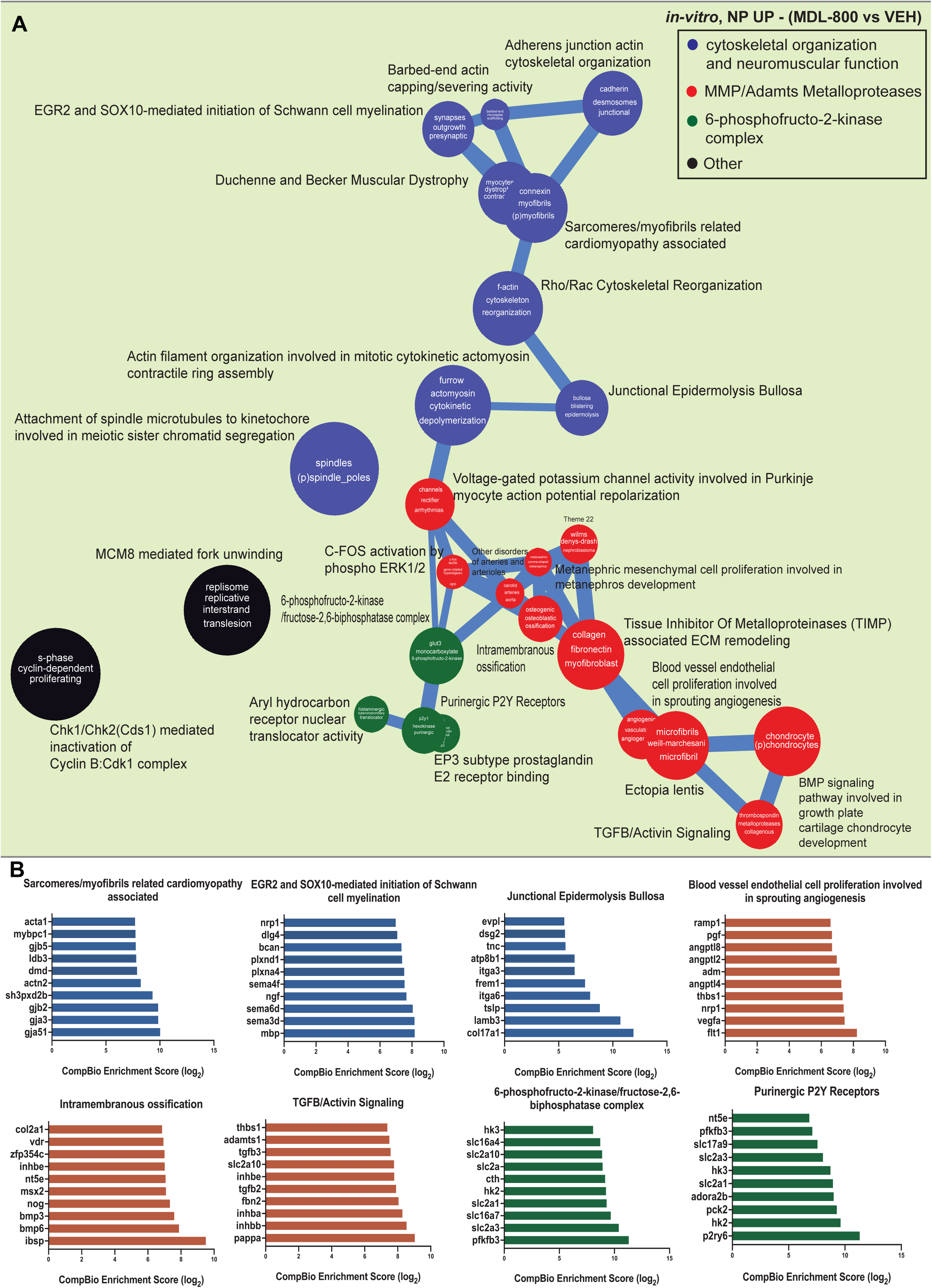
SIRT6 activation in NP cells results in clustering of upregulated DEGs in themes related to the cytoskeleton, cell junctions, skeletal tissues, extracellular matrix, and purinergic signaling. (A) CompBio analysis generated a thematic map of upregulated DEGs from an RNA sequencing experiment (Log_2_FC ≥1.5, FDR < 0.05) comparing MDL-800-treated vs VEH-treated NP cells, represented as a ball-and-stick model. Themes of interest are colored, and superclusters comprised of related themes are highlighted. (B) Top thematic DEGs plotted based on CompBio entity enrichment score. n = 4 independent experiments.

**Figure 9.**
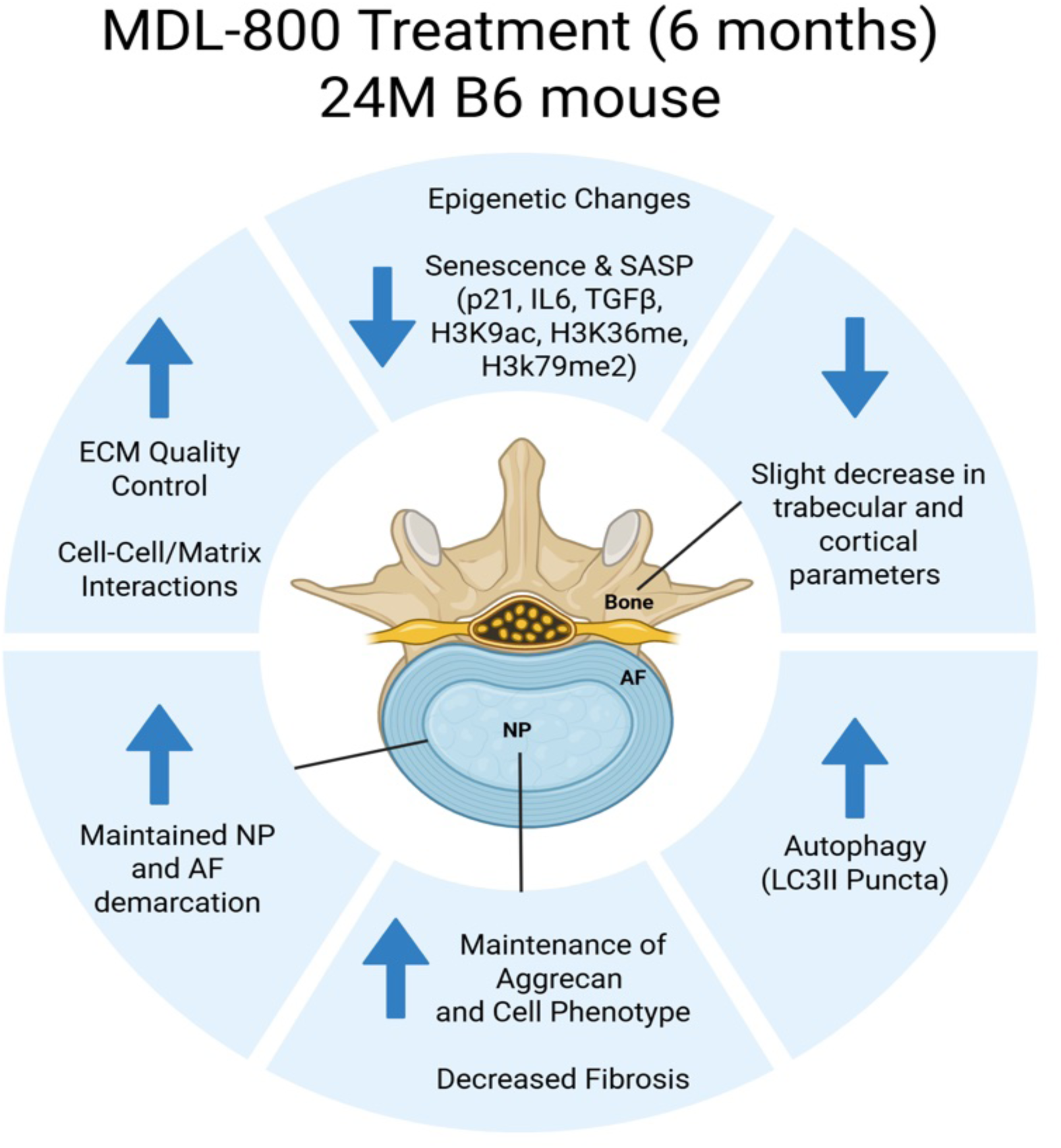
SIRT6 activation improves intervertebral disc health in the aging spine. A cartoon showing the major effects of SIRT6 activation on the intervertebral disc. SIRT6 activation by MDL-800 in old B6 mice diminishes intervertebral disc cell senescence, increases autophagy, and matrix homeostasis via epigenetic changes.

Next, using ATAC-seq data, we identified chromatin accessibility peaks and performed differential chromatin accessibility analysis between the groups. Chromatin accessibility signal profiles for each sample were visualized as heatmaps, and the average relative distribution of ATAC-seq peaks assigned to genomic features, including promoters, introns, exons, and distal regions, in vehicle control and MDL-800-treated NP cells was computed (Fig. S15A, B). We identified 251 upregulated and 21 downregulated shared genes between ATAC- and RNA-seq datasets (FC ≥ 2, FDR < 0.05), which are shown in a heatmap (top genes sorted based on ATAC-seq fold-changes) and a quadrant plot (Fig. S15C, D), demonstrating a strong correlation between chromatin accessibility and gene expression. Moreover, a set of seven genes *Serpine1, Evpl, Kcnh2, Gadd45g, Dlk2, Csrnp1,* and *Ptk7* were identified as consistently upregulated by MDL-800 treatment in both *in vivo* and *in vitro* NP RNA-seq datasets and shared with ATAC-seq data. *Serpine1 and Evpl are* shown as average chromatin accessibility maps (Fig. S15E). No genes were consistently downregulated across all three datasets.

### MDL-800 treatment results in vertebral bone changes and causes mild hyperalgesia, without mechanical allodynia, axial discomfort, or impairment of ambulation in mice

While there were no discernable differences in EP morphology between treatment groups (Fig. 1c), in the MDL-800 treated cohort, we noted a slight decrease in vertebral trabecular bone structural parameters (Fig. S16A), including BV/TV, trabecular number (Tb.N), and thickness (Tb. Th), without changes in trabecular spacing (Fig. S16B-E), suggesting a minimal effect on bone turnover. MDL-800 treated mice also exhibited a slight decrease in cortical bone parameters, including % closed porosity, cortical thickness (Cs. Th), again, notably, without changes in bone perimeter (B.Pm) (Fig. S16F-I). While these measurements suggested a small effect on vertebrae following long-term MDL-800 treatment, the effect size was too small to cause vertebral fractures. Notably higher integral and trabecular lumbar bone mineral density has been correlated to an increased risk of disc height loss as well as diminished disc area in humans (*41, 42*) suggesting a small decrease in bone structural parameters is likely to be protective.

To assess if MDL-800 treatment caused any behavioral changes, mechanical (Von Frey filament) and temperature (Hargreaves thermal; cold plantar assay) sensitization were measured in MDL- and vehicle-treated mice (Fig.S17A) (*43*). While MDL treatment did not alter the threshold to mechanical stimulation, we observed decreased latencies to cold and hot stimuli compared with vehicle controls (Fig.S17B-E). The open field test, however, showed comparable performance between cohorts, suggesting no adverse effect on overall ambulation in MDL-800 treated mice (Fig.S17F). Likewise, in both treatment cohorts, we observed a similar response to the grip strength test, an important metric used to assess frailty in humans and to measure axial pain in mice (Fig. S17G) (*44, 45*). To further study the locomotion-related parameters, gait analysis was performed. We did not observe changes in swing-, stance-, and stride durations, as well as stride length and frequency, suggesting the absence of impaired mobility or gait asymmetry (Fig. S17H-K).

## Discussion

Studies have demonstrated a strong link between SIRT6 activity and organismal aging, with *Sirt6* showing a significant correlation with increased lifespan in humans (*8, 46*), mice (*47, 48*), and other long-lived species (*49*). Notably, SIRT6 plays a prominent role in skeletal health (*19, 50*). Our previous loss-of-function studies have established a direct correlation between SIRT6 and intervertebral disc health in the aging spine (*51*). Disc cells reside in an avascular and therefore diffusively limited hypoxic niche and, consequently, possess limited regenerative capacity when they experience injury (*52*). Increased apoptosis, senescence, altered matrix synthesis, and activation or inhibition of catabolic processes involved in ECM turnover play pivotal roles during aging and degenerative conditions affecting the disc (*31, 53, 54*). While an earlier study showed suppression of cell senescence in an acute, traumatic disc injury model following SIRT6 activation (*26*), whether this approach will promote healthy spinal aging remains unknown. Notably, small-molecular SIRT6 activators have shown favorable outcomes in different preclinical disease models (*27–30*), underscoring their utility for *in vivo* interventions. Here, we show, for the first time, that systemic and long-term treatment of aged mice with MDL-800, a SIRT6 agonist, significantly improves outcomes in age-dependent disc degeneration and promotes healthier phenotypes. Mechanistically, SIRT6 activation results in diminished H3K9 acetylation along with H3K36 and H3K79 methylation. Furthermore, we noted a decreased senescence and SASP burden and increased autophagy in disc cells. This study not only establishes a positive link between SIRT6 activity and intervertebral disc health but also identifies mechanisms central to SIRT6 action on the disc.

Although SIRT6 regulates multiple signaling pathways and cellular processes across different tissues, many of its functions are tissue-type specific. Importantly, the enzymatic activity of SIRT6 - a class III histone deacetylase, long-chain fatty-acylase, and mono-ADP-ribosyl-transferase - depends on the availability of an essential cofactor, such as NAD⁺. Notably, NAD⁺ levels decline significantly with aging (*55*). Therefore, it is logical to conclude that reduced NAD⁺ availability would largely determine its enzymatic activity and functional impact in the aging spine. To overcome this challenge, we chose MDL-800, a highly selective allosteric activator of SIRT6, which, by directly binding to a surface allosteric site, enhances SIRT6’s affinity for both NAD+ and acylated substrates, thereby promoting catalytic efficiency (*56*). In this study, MDL-800 delivery to the disc and increased SIRT6 activation were evident by decreased H3K9 acetylation in NP cells.

SIRT6 plays a prominent role in the skeletal system (*57*). Particularly, loss of SIRT6 function in the intervertebral disc and knee articular cartilage leads to disc degeneration and osteoarthritis (*50, 51*). In line with these findings, Thompson grading in MDL-800 treated mice showed lower disc degeneration grades, suggesting improved histological outcomes of disc aging. Moreover, SIRT6-deficient NP cells and articular chondrocytes exhibited diminished antioxidant defense, increased senescence, DNA damage, and cell death (*51, 58*). Accordingly, decreased TUNEL-positive cells and higher GLUT1 and ACAN levels, indicating diminished cell death and maintenance of the NP cell phenotype, supported a positive role for SIRT6 in maintaining a healthy disc cell population.

In agreement with our previous observation of increased H3K9ac with methylation levels of H3K36 in Sirt6-deficient disc cells (*51*), SIRT6 activation resulted in diminished acetylation of H3K9, along with decreased methylation of H3K36 and H3K79. These studies also validated that SIRT6 does not regulate the acetylation status of H3K18 and H3K56 in the disc. Notably, studies have shown a positive correlation between cell senescence and H3K36me and H3K79me levels (*59–61*). Indeed, a significant decrease in p21 levels, as well as in important pro-fibrotic SASP molecules IL-6 and TGF-β, in MDL-800 treated mice highlighted that SIRT6 activation counteracts the senescence pathway. Accordingly, these context-specific roles of SIRT6 and its epigenetic functions underscore its importance in maintaining disc homeostasis during aging. Furthermore, we noted an overlap in enriched themes between the *in vivo* and *in vitro* transcriptomic datasets with select upregulated themes related to collagens, ECM turnover, cell-cell and cell-matrix adhesion, activin/BMP/FGF signaling, processes linked to disc health (*31, 53, 54*). On the other hand, downregulated shared themes were also linked to ECM, fibronectin, cytokines, and coagulation and clotting (fibrinogen, fibrinolysis, and plasmin). Notably, one of the shared upregulated genes, SERPIN1/PAI-1, reflects a broader decrease in the coagulation-related activity and improved phenotypic outcomes in discs of MDL-treated mice, as elevated fibrinolysis and plasmin activity are linked to the pathogenesis of OA (*62*). Supporting transcriptomic findings related to ECM homeostasis, we observed changes in the proportions of thin and thick collagen fibers in the AF, suggesting increased matrix turnover and a decreased incidence of NP fibrosis. Notably, these observations in concert with decreased SASP further supported the ability of SIRT6 to promote matrix homeostasis in the disc, as elevated TGF-β and profibrotic myofibroblast activation have been observed in models of Sirt6 deficiency, linking diminished SIRT6 to fibrosis in multiple tissues, including disc (*28, 29, 51*). Our results also align with the information-theoretic framework of aging, which characterizes aging as a function of epigenetic changes (*63*).

Autophagy is essential for maintaining physiological functions and cell survival in specialized tissue niches such as the disc (*2*). Noteworthy, one of the commonly dysregulated pathways during disc degeneration is autophagy. Indeed, in disc- and cartilage-targeted *Sirt6*^cKO^ mice, we noted decreased LC3 levels and a concomitant increase in the abundance of senescence markers (*51*). Notably, in MDL800-treated NP cells, we observed a marked increase in LC3II levels without affecting cellular bioenergetics, suggesting an important role for SIRT6 in promoting autophagy and inhibiting senescence (*64*). These findings are in line with a report showing that SIRT6 overexpression rescues acute injury-induced disc degeneration by modulating autophagy and senescence (*26*).

Since MDL800 was administered systemically, we investigated whether *Sirt6* activation alters the immune system and vertebrae. Notably, SIRT6 has been shown to modulate mast cell function (*65*) and macrophage polarization (*66*) and to suppress NF-κB-driven inflammation by enhancing IκBα expression and silencing pro-inflammatory gene expression (*66*). While we observed few changes in the immune cell populations in the spleen and bone marrow, enrichment of differentially expressed genes suggested a lower activation status of B cells and granulocytes. Notably, a recent study showed that B cell-deficient mice maintain a CD4 T cell compartment that resists immunosenescence and age-related decline in health (*67*). Accordingly, a decrease in B cell numbers in MDL800-treated mice and activation of defense response–related pathways in T cells and some decrease in B and T cell activation functionality in granulocytes likely suggest a favorable and non-activated immune cell profile. Importantly, stable plasma cytokine and broader metabolomes, and the lack of alterations in inflammatory marker GlyA, suggested no adverse effects on the immune system and metabolic health. A small decrease in a select bone structural parameter was noted and may have resulted from SIRT6’s ability to promote osteoclast differentiation from hematopoietic precursors, but a relatively smaller stimulatory effect on osteoblast function (*68*). However, the absence of adverse events, such as vertebral fractures, the lack of changes in overall ambulation, gait, and axial pain, and the known inverse correlation between lumbar trabecular bone mineral density and the risk of disc degeneration in humans (*41, 42*), suggest a rather protective adaptive change. It must also be acknowledged here that while we only studied male mice due to the known robust effects of SIRT6 on multiple age-related phenotypes in males, future work should decipher the role of SIRT6 activation on spinal health of females.

In summary, our studies demonstrate a positive relationship between SIRT6 and intervertebral disc health in vivo. SIRT6 activation, by inhibiting cell death and senescence, promoting autophagy, and maintaining ECM homeostasis, contributed to healthier discs in MDL800-treated mice. Modulating SIRT6 activity through pharmacological activation with specific, well-tolerated agonists offers a non-invasive, translational strategy to ameliorate age-dependent disc degeneration and preserve disc health in the aging spine.

## Materials and Methods

### Mouse Studies

Animal studies were approved by the Institutional Animal Care and Use Committee of Thomas Jefferson University following guidelines from the National Institutes of Health Guide for the Care and Use of Laboratory Animals. 18-month-old male C57BL/6 mice (NIA colony, Charles River) were housed in groups of 4 per cage and had access to water and food ad libitum. Mice were randomly divided into two groups, and except for 2 doses/week in the first month, received weekly intraperitoneal injections of either DMSO (Vehicle Control group) or 75 mg/kg of MDL-800 (SIRT6 activator) in DMSO for six months, i.e., up to 24 months of age, n=13 Control, 15 MDL-800 mice. We chose to use male mice based on our recent study, which showed enhanced disc degeneration in male C57BL/6J mice following SIRT6 deletion, as well as previous studies demonstrating larger effect sizes on lifespan and protective effects against age-associated changes in male mice (*12, 51*).

### Behavioral studies

Mice were acclimated to the behavioral testing room for 1 hour before all behavioral tests. Forelimb grip strength to measure the frailty and axial pain, open field test to assess the general locomotion, Von Frey test for mechanical allodynia, Hargreaves thermal and cold plantar assays to measure hyperalgesia were performed (n = 7 Vehicle Control, n = 9 MDL mice) (*43, 70*). Gate analysis was performed to understand any changes in the ambulation (*43*) (see Suppl. Methods for details)

### Histological studies

Spines were dissected and fixed in 4% paraformaldehyde (PFA) for either 6 hours or 48 hours, followed by decalcification in 20% EDTA at 4°C before embedding in OCT or paraffin for sectioning. 7μm midcoronal sections from four lumbar levels (L3-S1) of each mouse (n = 7 Vehicle Control, n = 9 MDL mice; 2-4 discs/mouse) were stained with Safranin-O/Fast Green/Hematoxylin for histological assessment using a modified Thompson grading scale for NP and AF by at least three blind observers. Similarly, Picrosirius Red staining was performed to characterize collagen fibers in tissue sections. Safranin-O staining was visualized using an Axio Imager 2 microscope (Carl Zeiss, Germany) using 5×/0.15 N-Achroplan or 20×/0.5 EC Plan-Neofluar objectives and Zen2TM software (Carl Zeiss).The heterogeneity of collagen organization was evaluated using a polarizing light microscope, Eclipse LV100 POL (Nikon, Tokyo, Japan), equipped with a 10x/0.25 Pol/WD 7.0 objective and a DS-Fi2 camera. Images were analyzed using NIS Elements AR 4.50.00 (Nikon, Tokyo, Japan). Under polarized light, as the thickness of collagen fibers increases, their birefringence color changes from green to yellow to orange to red, *i*.*e*., from shorter to longer wavelengths(*71, 72*), allowing quantitative assessment of compositional information in tissue sections. Color threshold levels were maintained constant between all analyzed images.

### Immunohistological analyses

Deparaffinized sections following antigen retrieval or frozen sections were blocked in 5% normal serum in PBS-T, and incubated with antibodies against H3K9Ac (1:50, Sigma, 06-942), Collagen I (1:100, Abcam ab34710), Collagen X (1:500, Abcam ab58632), Aggrecan (1:300, Abcam ab11570); GLUT1 (1:200, Abcam, ab40084), F-CHP (1:100, 3-Helix), p21 (1:200, Novus NB100-1941), IL-6 (1:50, Novus NB600-1131), TGF-ý (1:200, Abcam; ab92486) (n= 4-6 mice/group, 2-3 discs/mouse were analyzed). For mouse antibodies, a MOM kit (Vector Laboratories, BMK-2202) was used for blocking and primary antibody incubation. Tissue sections were washed and incubated with species-appropriate Alexa Fluor-594 conjugated secondary antibodies (Jackson ImmunoResearch,1:700). TUNEL staining was performed using the *In situ* cell death detection kit (Roche Diagnostics). Briefly, sections were deparaffinized and permeabilized using Proteinase K (20 μg/mL), and the TUNEL assay was performed according to the manufacturer’s protocol. The sections were mounted with ProLong Gold Antifade Mountant with DAPI (Fisher Scientific, P36934), visualized using an Axio Imager 2 microscope with 5×/0.15 N-Achroplan or 20×/0.5 EC Plan-Neofluar objectives, and images were captured with an Axiocam MRm monochromatic camera (Carl Zeiss) and Zen2 software (Carl Zeiss AG, Germany). Both caudal and lumbar discs were used for the analysis. Staining area and cell number quantification were performed using the ImageJ software, v1.53e, (http://rsb.info.nih.gov/ij/).

### Disc tissue RNA-sequencing

NP and AF tissues were dissected from Vehicle control and MDL-800-treated mice (n = 5 mice/group). Samples were homogenized, and DNA-free, total RNA was extracted using the RNeasy® Mini kit (Qiagen). RNA with RIN > 7 was used for RNA-sequencing analysis. Significantly up- and downregulated DEGs (FC> 1.5, p< 0.05) were analyzed using the GTAC-CompBio Analysis Tool (https://gtac-compbio-ex.wustl.edu). CompBio, a multiomic analysis platform built on an episodic and semantic memory-based intelligence engine, is used to analyze DEGs. This engine stores PubMed abstracts and full-text articles as memories, referencing the input DEGs to identify relevant processes and pathways. Conditional probability analysis is used to calculate the statistical enrichment score for biological concepts (processes/pathways) relative to those expected by random sampling. The scores are then normalized for significance empirically over a large, randomized query group. Related concepts derived from DEGs are grouped into broader, higher-level themes, which represent overarching biological patterns, such as pathways, processes, cell types, and structures as described previously(*73, 74*).

### ScRNA-seqencing of splenocytes and bone marrow cells

Spleens were collected from Vehicle Control and MDL-treated mice. Splenocytes were dissociated from the tissue matrix by gentle compression through a 100-μm strainer (StemCell Technologies, 27217). Similarly, bone marrow cells were isolated by gently flushing the femur and humerus. Thereafter, RBC lysis (BioLegend, 420302) was performed, followed by two PBS + 3% FBS washes. ScRNA-sequencing was performed on freshly isolated cells using the standard 10X protocol (see Supplemental Methods). Raw sequencing data were aligned to the mouse reference transcriptome (mm10-2020-A) and processed with Cell Ranger Multi version 7.0.1. The count matrix was analyzed using Seurat version 4.3.0. Briefly, data were converted into Seurat Objects using the CreateSeuratObject function, and all cells were analyzed for their unique molecular identifier and mitochondrial gene fractions; cells were included if 200 < *n*feat < 2500, and mitochondrial genes accounted for <5%. Data were integrated with logarithmic normalization (LogNormalize) and organized according to variation in the data using FindVariableFeatures. Dimensional reduction was performed using principal components analysis, and the FindNeighbors (dims = 1:20) function, followed by the FindClusters function, was used to calculate the number of cell clusters using the Louvain algorithm, with the resolution parameter set to 0.5. DimPlot was then used to visualize the data as UMAP. Cell types for each condition were determined using SingleR, with the MouseRNAseqData reference dataset and main labels. The clusterProfiler package was used to conduct GSEA on DEGs (FDR < 0.05) identified in various tissue and treatment-based comparisons.

### Plasma Cytokine analysis

Blood was collected by intracardiac puncture in heparin-coated syringes and centrifuged at 1500 relative centrifugal force (rcf) at 4°C for 15 min to isolate the plasma (n =10 mice/group). Samples were stored at −80°C until analysis. According to the manufacturer’s specifications, levels of pro-inflammatory proteins, cytokines, and TH17 mediators were analyzed using the V-PLEX Mouse Cytokine 29-Plex Kit (Meso Scale Diagnostics, Rockville, MD).

### Plasma metabolomics by LC/MS and NMR

Plasma LC/MS analysis was performed as described previously (n =8 Vehicle, 7 MDL mice) (75). Briefly, polar metabolites were extracted from 50 µl plasma samples with 500 µl ice-cold 80% methanol, and deproteinated supernatants were stored at −80 °C prior to analysis. A quality control (QC) sample was generated by pooling equal volumes of all samples after extraction. LC-MS analysis was performed on a Thermo Scientific Q-Exactive HF-X mass spectrometer with HESI II probe and Vanquish Horizon UHPLC system. Hydrophilic interaction liquid chromatography (HILIC) was performed at a flow rate of 0.2 ml/min on a ZIC-pHILIC column (150 × 2.1 mm, 5 µm particle size, EMD Millipore) with a ZIC-pHILIC guard column (20 × 2.1 mm, EMD Millipore) at 45 °C. Solvent A was 20 mM ammonium carbonate, 0.1% ammonium hydroxide, pH 9.2, and solvent B was acetonitrile. The gradient was 85% B for 2 min, 85% B to 20% B over 15 min, 20% B to 85% B over 0.1 min, and 85% B for 8.9 min. The autosampler was held at 4 °C. For each analysis, 4 µl of the sample was injected. The following parameters were used for the MS analysis: sheath gas flow rate, 40; auxiliary gas flow rate, 10; sweep gas flow rate, 2; auxiliary gas heater temperature, 350 °C; spray voltage, 3.5 kV for positive mode and 3.2 kV for negative mode; capillary temperature, 325 °C; and funnel RF level, 40. All samples were analyzed by full MS with polarity switching. The QC sample was analyzed at the start of the sample sequence and after every 8–14 samples. The QC sample was also analyzed by data-dependent MS/MS with separate runs for positive and negative ion modes. Full MS scans were acquired at 120,000 resolution with an automatic gain control (AGC) target of 1e6, maximum injection time (IT) of 100 ms, and scan range of 65–975 m/z. Data-dependent MS/MS scans were acquired for the top 10 highest intensity ions at 15,000 resolution with an AGC target of 5e4, maximum IT of 50 ms, isolation width of 1.0 m/z, and stepped normalized collision energy (NCE) of 20, 40, 60. Data analysis was performed using Compound Discoverer 3.1 (ThermoFisher Scientific) with separate analyses for positive and negative polarities. Results were manually processed to remove entries with apparent peak misintegrations and correct commonly misannotated metabolites. Positive and negative data sets of identified compounds were merged, and the preferred polarity was selected for compounds identified in both polarities. Compound quantifications were normalized per volume of plasma injected, which was equivalent for all samples. Values from the study were further normalized to the summed area of identified metabolites in each sample. For compounds identified multiple times at different retention times, a single entry was selected with priority given to standards database matches, followed by greater mzCloud match factors and peak areas.

Additionally, compounds shown in Figures S2 and S3 were measured from plasma samples (n =8 Vehicle Control, 7 MDL mice) at Labcorp (Labcorp, Morrisville, NC). NMR spectra were acquired on a 400 MHz NMR instrument, the Vantera Clinical Analyzer, from EDTA plasma samples as described for the *NMR LipoProfile* test. The data were transferred and processed using the LP4 lipoprotein profile deconvolution algorithm, as reported in the supplemental methods (*76, 77*).

### Micro-CT analysis

Micro-CT (μCT) scanning (Bruker SkyScan 1275) was performed on fixed spines using parameters of 50 kV (voltage) and 200 μA (current) at 15 μm resolution as previously described (n =8 Vehicle Control, 12 MDL mice, 4 vertebrae/mouse) (31, 51). Images were reconstructed using the nRecon program (Version: 1.7.1.0, Bruker) and analysis was performed using CTan (version 1.17.7.2, Bruker). Disc height and vertebral length were measured at three different points equidistant from the center of the bone on the sagittal plane and used to calculate the Disc height index (DHI).

### NP cell isolation and treatments

Primary NP cells from adult Sprague Dawley rats (3-6-month-old, Charles River) were isolated and cultured in antibiotic-supplemented complete DMEM containing 5 mM glucose and 10% FBS. To explore the role of SIRT6 *in vitro*, cells were treated with either DMSO (1:2000) or MDL-800 (25 μM in DMSO) in 1% hypoxia for 24-96 hours.

### Immunoblotting

Vehicle control and MDL-800-treated (25 μM) NP cells (n = 4 independent experiments) were used to extract either histones (Abcam-ab113476) or total cellular proteins. 10-40 μg of protein was electroblotted to nitrocellulose membranes (Amersham, GE, Burlington, MA, USA). The membranes were blocked and incubated overnight at 4°C with antibodies against H3K9ac (1:1000, 06-942S, Millipore), Histone 3 (1:1000, 9715), SIRT6 (1:1000, D8D12-12486), LC3 (1:1000, 12741), GAPDH (1:3000, 5174) from Cell Signaling Technology. Immunolabeling was detected on the Azure 300 system using an ECL reagent (Azure Biosystems, Dublin, CA), and densitometric analysis was performed with ImageJ.

### Histone ELISA

The Histone Modification Multiplex Assay kit (ab185910, Abcam) was used to quantify Histone 3 modifications, including lysine acetylation and mono -, di-, and tri-methylation, using histones isolated from NP cells treated with Vehicle control and MDL-800 for 24 hours, according to the manufacturer’s instructions (n = 3 independent experiments).

### NP cell RNA- and ATAC-Sequencing

Following vehicle control and MDL-800 treatment (25 μM) of rat NP cells for 24 hours, total DNA-free RNA was extracted using RNeasy mini columns (Qiagen) (n=4 independent experiments). The extracted RNA with RIN > 7 was used for RNA sequencing, and analysis of DEGs (Log_2_FC ≥1.5, FDR≤0.05) was performed with the GTAC-CompBio tool (https://gtac-compbio-ex.wustl.edu). For ATAC-sequencing, Vehicle control and MDL-800-treated NP cells were trypsinized and collected by centrifugation at 600-800g for 5 min at 4°C. Cells were resuspended in 500 μL cryopreservation medium containing 50% serum and 10% DMSO and then cryopreserved in 2 mL cryopreservation tubes. The tubes were shipped to Azenta for sequencing using standard protocols. Briefly, cells were thawed and treated with Tn5 transposase, and tagmented DNA fragments were purified using a DNA purification kit for library synthesis (Azenta). Purified sequencing libraries were used for paired-end 150-bp-long reads on Illumina NovaSeq. Reads were mapped to the reference genome with default parameters. Aligned ATAC-seq reads were used to remove duplicates, and peak calling was performed, followed by a multi-omic analysis of ATAC-seq and RNA-seq (FC≥2, FDR≤0.05) (see Supplemental Methods).

### Statistics

All statistical analyses were performed using Prism7 or above (GraphPad, La Jolla). Since the unique interactions between genetic, biological, and biomechanical factors at individual spinal levels have shown to produce different phenotypic outcomes, each disc was considered as an independent sample (*31, 78, 79*). Data are represented as box and whisker plots with median, minimum, and maximum values or mean ± SEM. Data distribution was assessed using normality tests, and differences between the two groups were analyzed using an unpaired t-test. The differences between the three groups were analyzed by ANOVA or Kruskal–Wallis for non-normally distributed data. A chi-square (χ2) or Fisher test, as appropriate, was used to analyze the differences between the distribution of percentages. p ≤ 0.05 was considered a statistically significant difference.

## Supporting information

CompBio Enrichment list

## Acknowledgments

This study was supported by the Michael Michelson Gift Fund, the NIA grant R01AG073349, and the NIAMS grant R01AR055655 (M.V.R.). We thank the Proteomics and Metabolomics Core Facility at Wistar Institute for LC-MS analysis. We thank Victoria Tran, Tanvi Abhang, Cindy Pham, and Esther Akande for technical help.

## Author Contributions

M.V.R and J.C. conceptualized the study, P.R. and M.V.R. designed the experiments. P.R., B.W., S.J., S.E.B., M.T. and Q.W. performed the experiments, collected, and analyzed the data. O.O, R.A.B., O.A., and R.M. performed bioinformatics analysis. P.R. and M.V.R. interpreted the results, and M.V.R. wrote the original draft of the manuscript. All authors reviewed and approved the final draft of the manuscript.

## CONFLICT OF INTERESTS

Q.W. is an employee of Labcorp. The authors of this manuscript do not have conflicts of interest to disclose.

## DATA AVAILABILITY

Sequencing data associated with this study are deposited in the GEO database under accession GSE320197 and GSE320429. All datasets generated and analyzed during this study are included in this published article.

## Disclosures

None

## ETHICS STATEMENT

All animal experiments were performed in accordance with protocols approved by the Thomas Jefferson University IACUC.

## Notes

### Competing Interest Statement

The authors have declared no competing interest.

