## Supplementary Methods for "SIRT6 Activation Improves Intervertebral Disc Health in the Aging Spine"

**Epigenetic Activation of SIRT6 Protects the Aging Intervertebral Disc**

**Supplemental Methods**

**Behavioral Studies**

Mice (n = 7 Vehicle Control,  9 MDL mice) were acclimatized to the behavior testing room for 1 hour before testing for all behavior tests. Forelimb grip strength was assessed using a Grip Strength Meter (DFIS-2 Series Digital Force Gauge, Columbus Instruments). To measure grip strength, animals were gently held by their tails and were allowed to tightly grasp a force gauge bar using both forepaws. Mice were then pulled away from the gauge until both limbs released the bar. The data were recorded and are representative of the average of five trials per mouse. Between trials, mice were rested for sixty seconds. An open field test was used to assess the general locomotion of mice. Mice were placed in an open field apparatus and recorded with an overhead camera for 10 mins. Data from these videos was processed in MATLAB using the open-source code developed by Zhang et al. (1) to determine the distance traveled by each mouse.

Modified versions of the Hargreaves thermal and cold plantar tests were used to assess thermal hyperalgesia mice. Mice were gently held by the scruff as their hind paw was placed over either an infrared heart source or a piece of dry ice covered by a glass plate. The time taken for the mouse to withdraw its paw from the stimulus was recorded. The trials were done in triplicate with a 2 min. rest between each. There was a 30s cut off for exposure to each thermal stimulus.

A modified version of the von Frey filament test was conducted to assess tactile sensitivity. Mice were placed in a glass box with a mesh bottom, sized so that movement was limited but not fully prevented. A series of filaments, producing between .6 and 2 g of force, were applied to the center of each hind paw. Paw withdrawal by the mice led to the next filament of a lesser force being applied, while no response led to the next filament of a stronger force being applied. This was done for 10 trials on each foot. The filament that led to a paw withdrawal at least half of the time is defined as the withdrawal threshold.

Gait analysis was performed to assess pain associated with locomotion. Mice were individually placed on a transparent treadmill belt, encased within a plexiglass box. After 2 minutes of acclimation, the treadmill was set to 8 cm/s. DigiGait’s ventral plane imaging technology captured and analyzed various gait features, including stride, swing, and stance.

**ScRNAseq**

Bone marrow and splenocyte samples were uniquely barcoded using the 10x Genomics 3’ CellPlex Kit (10x Genomics, Pleasanton, CA) as per manufacturer’s instructions. A total of 4 samples of each sample type were multiplexed together into a single lane on the Chromium Controller (10x Genomics), for targeted capture of 2,500 single cells/sample. Two 10x lanes were loaded, one for bone marrow and the other for splenocytes. Single cell droplets were generated using the Chromium Next GEM Single Cell 3’ Reagent Kits v3.1 (Dual Index) with Feature Barcode technology for Cell Multiplexing (10x Genomics). The following PCR cycles were used at each step: 11 PCR cycles for cDNA amplification, 6 PCR cycles for Cell Multiplexing Library Construction, 14 PCR cycles for index incorporation into final gene expression library. The overall library size was assessed using the 4200 TapeStation and the High-sensitivity DNA 5000 ScreenTape (Agilent, Santa Clara, CA).  Concentration was determined using the Qubit Fluorometer 2.0 (Thermofisher, Waltham, MA). 2 gene expression libraries and 2 cell multiplexing libraries were pooled according to the following reads per cell requirement suggested by 10x Genomics: 20,000 reads/cell for gene expression analysis and 2,000 reads/cell for multiplexing barcode identification. Next Generation Sequencing on the pool was done on the NextSeq 2000 (Illumina, San Diego, CA) using a P2 100 cycle kit (Illumina) yielding 450 million total reads, paired end run with the following run parameters: 28 base pair x 10 base pair (index) x 90 base pair.

**LC/MS data analysis**

Retention time alignment used the adaptative curve model with 0.3 min maximum shift, 5 ppm mass tolerance, and 3 S/N threshold. Peak detection required less than 5 ppm mass error for extracted ion chromatograms with a 50,000 minimum peak intensity. [M + H] + 1 and [M-H]-1 were set as base ions with consideration for other adducts. Peaks were required to have a width at half height less than 0.5 min and a minimum of 5 scans. Components that had only a monoisotopic peak and no further isotopes were not considered. The maximum element count for isotope pattern modeling was C90H190N10Na2O15P3S5. Compounds were grouped across samples with 5 ppm mass error and 0.3 min retention time shift. Peaks not initially detected in a given sample were determined using the fill gaps algorithm with a 5 ppm mass error and a 1.5 S/N threshold for real peak detection. The gap function uses a priority system to determine missing values: (1) matching detected ions based on expected m/z and retention time regardless of adduct assignment, (2) re-detecting peaks at lower thresholds, (3) simulating peaks based on expected m/z, and (4) imputing spectrum noise based on detection limit values. Compound quantifications were corrected for instrument drift by QC areas using the cubic spline regression model. Each compound was required to be detected in at least 40% of QC runs with a Relative Standard Deviation (RSD) less than 50%. Metabolites were identified by accurate mass (5 ppm mass error) and retention time (0.5 min shift) using a database generated from pure standards or by accurate mass and MS2 spectra using the mzCloud spectral database (mzCloud.org), specifically the ‘Endogenous Metabolites’ and ‘Steroids/Vitamins/Hormones’ compound classes and selecting the best matches with HighChem HighRes identity search match factors of 50 or greater.

**NMR data parameters**

The *NMR MetaboProfile* analysis, using the LP4 lipoprotein profile deconvolution algorithm, reports lipoprotein particle concentrations and sizes, as well as concentrations of metabolites such as total branched-chain amino acids, valine, leucine, isoleucine, alanine, glucose, citrate, total ketone bodies, β-hydroxybutyrate, acetoacetate, and acetone (Labcorp, Morrisville, NC). The diameters of the various lipoprotein classes and subclasses are total triglyceride-rich lipoprotein particles (TRL-P) (24–240 nm), very large TRL-P (90–240 nm), large TRL-P (50–89 nm), medium TRL-P (37–49 nm), small TRL-P (30–36 nm), very small TRL-P (24–29 nm), total low-density lipoprotein particles (LDL-P) (19–23 nm), large LDL-P (21.5–23 nm), medium LDL-P (20.5–21.4 nm), small LDL-P (19–20.4 nm), total high-density lipoprotein particles (HDL-P) (7.4–13.0 nm), large HDL-P (10.3–13.0 nm), medium HDL-P (8.7–9.5 nm), and small HDL-P (7.4–7.8 nm). Mean TRL, LDL, and HDL particle sizes are weighted averages derived from the sum of the diameters of each of the subclasses multiplied by the relative mass percentage. Linear regression against serum lipids measured chemically in an apparently healthy study population (*n* = 698) provided the conversion factors to generate NMR-derived concentrations of total cholesterol (TC), triglycerides (TG), TRL-TG, TRL-C, LDL-C, and HDL-C. NMR-derived concentrations of these parameters are highly correlated (*r* ≥ 0.95) with those measured by standard chemistry methods. Details regarding the performance of the assays that quantify BCAA, alanine, and ketone bodies have been reported (*2, 3*).

**RNAseq-ATACseq multiomic analysis**

Bulk RNA-seq was generated by Azenta Life Sciences using Rat RNA libraries and sequenced on a NovaSeq platform (2 × 150 bp). Data quality was assessed for all samples. Raw reads were trimmed using Trimmomatic (*4*) tool (v0.36). The reads were then aligned to the Rat reference genome (Rnor_6.0, Ensembl) using STAR (*5*) (v2.5.2b). Gene quantification was handled using featureCounts (*6*) tool from the Subread package (v1.5.2), and DESeq2 (*7*) tool was considered for differential expression analysis. Significantly differentially expressed genes were selected using a threshold of FC ≥ 2, FDR ≤ 0.05. A volcano plot was generated using the ggplot2 R package (*8*). ATAC-seq libraries were sequenced on an Illumina HiSeq platform (2 × 150 bp). Adapter trimming and removal of low-quality bases were performed using Trimmomatic (*4*) (v0.38). Trimmed reads were aligned to the rat reference genome using Bowtie2 (*9*). Chromatin accessibility peaks were identified using MACS2 (*10*) (v2.1.2), and differential chromatin accessibility analysis was performed using DiffBind (*11*).

RNA-seq and ATAC-seq datasets were integrated by intersecting statistically significant (FC ≥ 2, FDR ≤ 0.05) gene lists from each modality, which identified 251 upregulated and 21 downregulated genes. Furthermore, a set of 7 genes was identified as consistently upregulated in the *in vitro* and *in vivo* MDL-800-treated RNA-seq and ATAC-seq samples compared to the vehicle. No genes were consistently downregulated across all datasets. The ATAC-seq peaks were shown using the Integrative Genomics Viewer (*12*) (IGV) software. Heatmaps showing chromatin accessibility signal profiles were generated using deepTools (*13*). Signal matrices were computed using the *computeMatrix scale-regions function* and visualized using *plotHeatmap*. For each condition, three biological replicates were visualized with average signal profiles shown above the heatmaps. The relative distribution of ATAC-seq peaks in the control and vehicle samples, annotated to genomic features (e.g., promoters, introns, exons, and distal regions). Additionally, a heatmap showing RNA-seq gene expression profiles (top 50 DEGs UP and 21 Down) was generated using the heatmaply (*14*) R package. A Quadrant plot was generated using ggplot2 (*8*) to visualize concordant and discordant changes in gene expression and chromatin accessibility.

**Seahorse XF analysis**

In brief, NP cells were plated in a 24-well Seahorse V7-PS test plate, treated with MDL-800 or Vehicle for 4 days, and placed in hypoxia (1% O_2_) 24 hours before the experiment (n = 4 independent experiments, 4 replicates/experiment). On the day of the experiment, cells were washed three times with 700 μL of KRPH (Krebs-Ringer phosphate HEPES) and incubated with KRPH + BSA for 1 hour at 37 °C. Seahorse XFe24 flux analyzer (Agilent Technologies) was used to determine maximum glycolytic capacity and ATP production rate using methods reported by Mookerjee et al. (*15*) Experimental design for ATP-consumption included sequential additions of 10 mM glucose, 1 μM rotenone plus 1 μM myxothiazol, 2 μg/mL oligomycin. To measure glycolytic capacity, sequential additions of 10 mM glucose, 1 μM rotenone plus 1 μM myxothiazol, and 200 μM monensin plus 1 μM FCCP were performed. The normalized traces for oxygen consumption rate (OCR) and extracellular acidification rate (ECAR) were used to calculate experimental parameters.
