## Supplementary Figures for "SIRT6 Activation Improves Intervertebral Disc Health in the Aging Spine"

**Figure S.1**

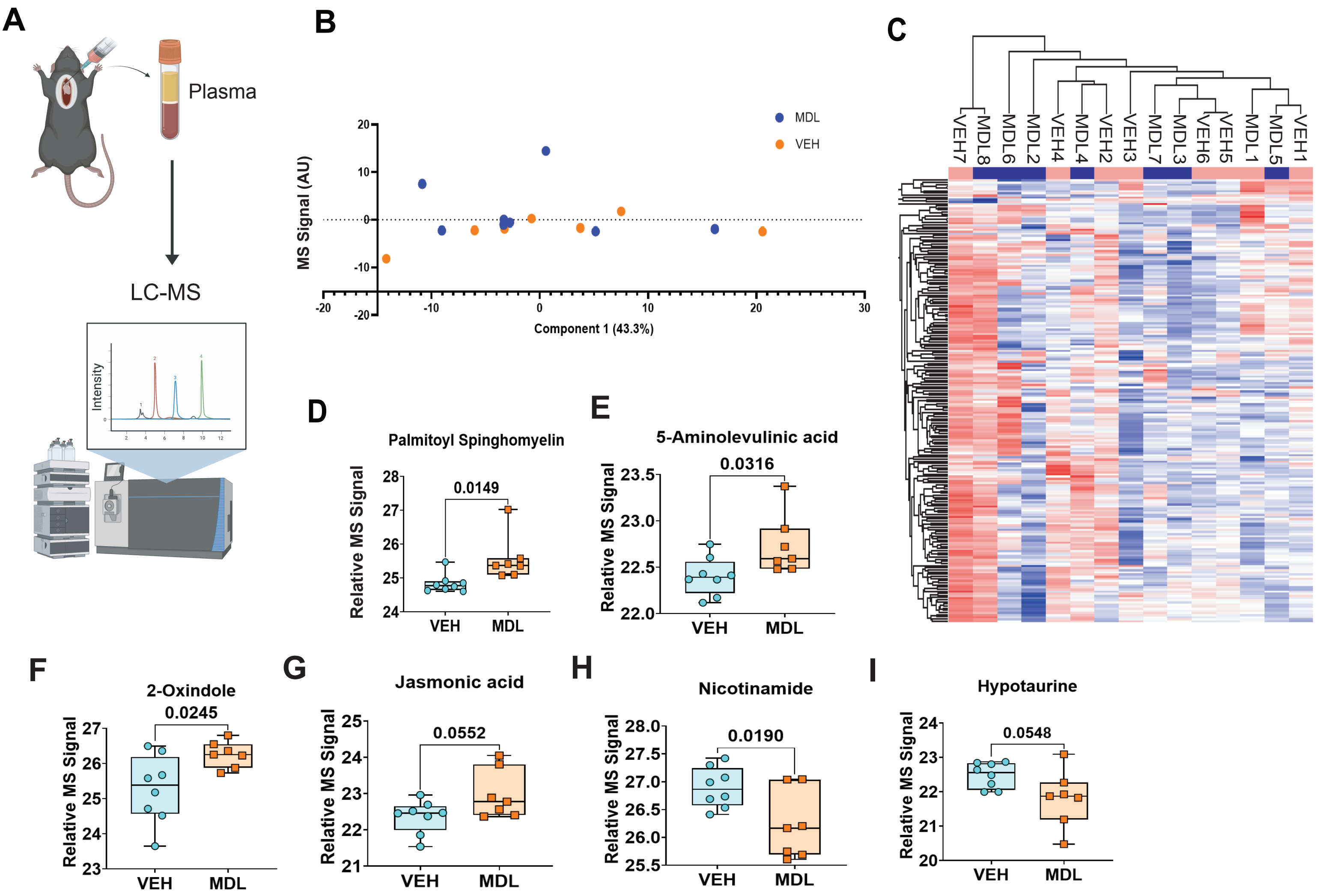

A
Plasma
LC-MS
Intensity
B
MS Signal (AU)
Component 1 (43.3%)
MDL
VEH
C
VEH7
MDL8
MDL6
MDL2
VEH4
MDL4
VEH2
VEH3
MDL7
MDL3
VEH6
VEH5
MDL1
MDL5
VEH1
D
Palmitoyl Spinghomyelin
0.0149
Relative MS Signal
VEH
MDL
E
5-Aminolevulinic acid
0.0316
Relative MS Signal
VEH
MDL
F
2-Oxindole
0.0245
Relative MS Signal
VEH
MDL
G
Jasmonic acid
0.0552
Relative MS Signal
VEH
MDL
H
Nicotinamide
0.0190
Relative MS Signal
VEH
MDL
I
Hypotaurine
0.0548
Relative MS Signal
VEH
MDL

Figure S.2

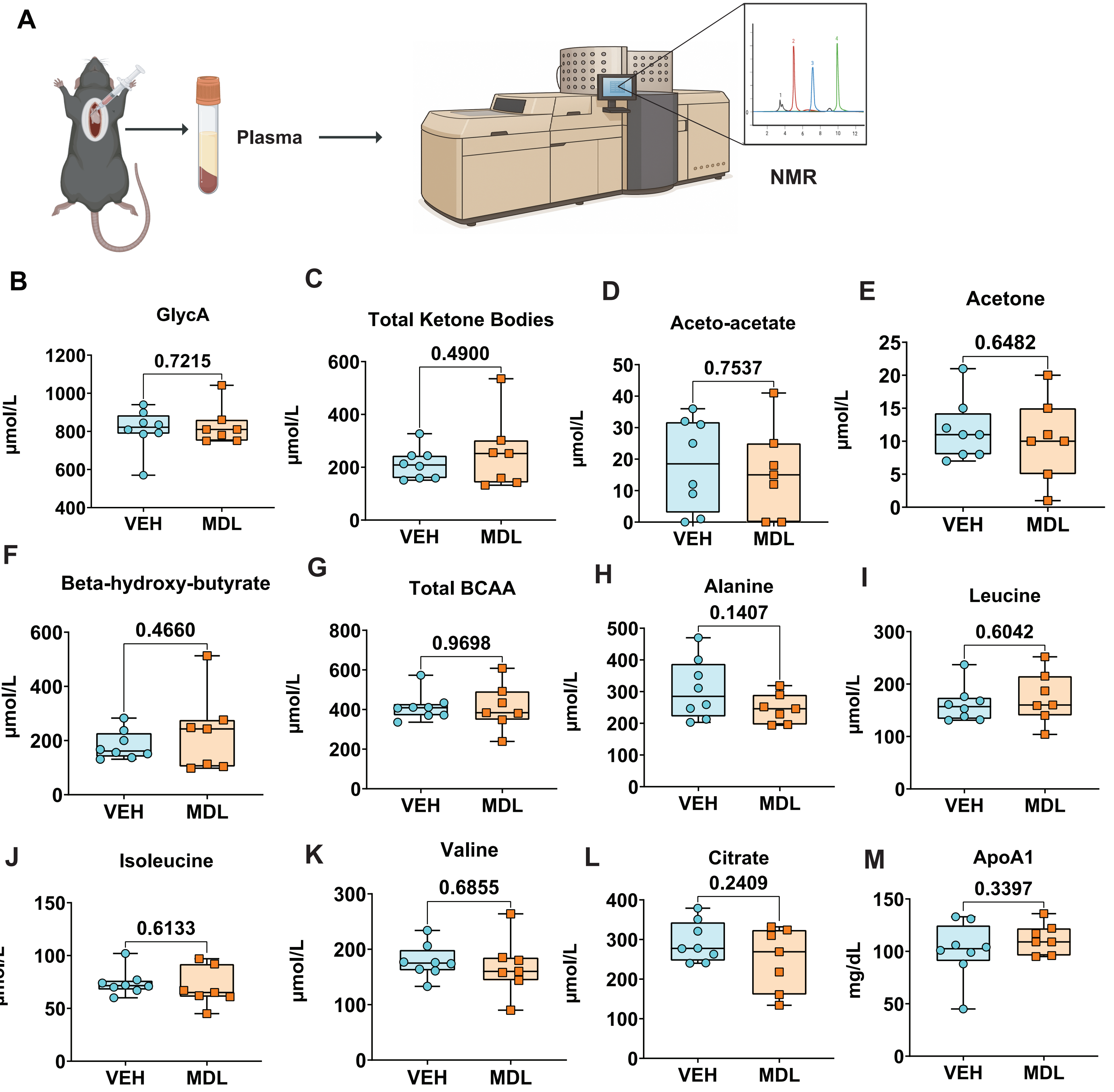
A
Plasma
NMR
B
GlycA
0.7215
μmol/L
VEH
MDL
C
Total Ketone Bodies
0.4900
μmol/L
VEH
MDL
D
Aceto-acetate
0.7537
μmol/L
VEH
MDL
E
Acetone
0.6482
μmol/L
VEH
MDL
F
Beta-hydroxy-butyrate
0.4660
μmol/L
VEH
MDL
G
Total BCAA
0.9698
μmol/L
VEH
MDL
H
Alanine
0.1407
μmol/L
VEH
MDL
I
Leucine
0.6042
μmol/L
VEH
MDL
J
Isoleucine
0.6133
μmol/L
VEH
MDL
K
Valine
0.6855
μmol/L
VEH
MDL
L
Citrate
0.2409
μmol/L
VEH
MDL
M
ApoA1
0.3397
mg/dL
VEH
MDL

Figure S.3

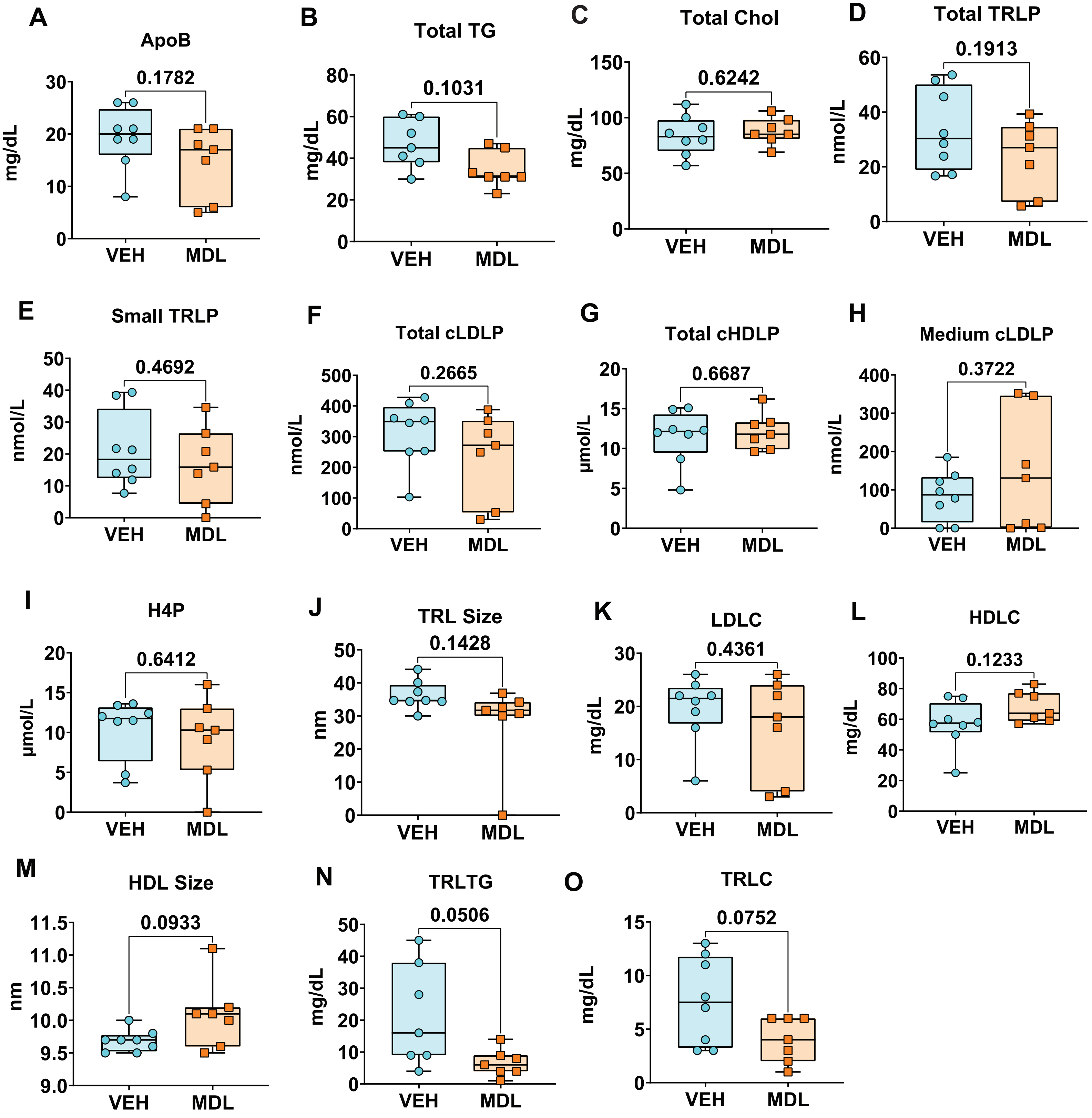
A
ApoB
0.1782
mg/dL
0
10
20
30
VEH
MDL
B
Total TG
0.1031
mg/dL
0
20
40
60
80
VEH
MDL
C
Total Chol
0.6242
mg/dL
0
50
100
150
VEH
MDL
D
Total TRLP
0.1913
nmol/L
0
20
40
60
VEH
MDL
E
Small TRLP
0.4692
nmol/L
0
10
20
30
40
50
VEH
MDL
F
Total cLDLP
0.2665
nmol/L
0
100
200
300
400
500
VEH
MDL
G
Total cHDLP
0.6687
µmol/L
0
5
10
15
20
VEH
MDL
H
Medium cLDLP
0.3722
nmol/L
0
100
200
300
400
VEH
MDL
I
H4P
0.6412
µmol/L
0
5
10
15
20
VEH
MDL
J
TRL Size
0.1428
nm
0
10
20
30
40
50
VEH
MDL
K
LDLC
0.4361
mg/dL
0
10
20
30
VEH
MDL
L
HDLC
0.1233
mg/dL
0
20
40
60
80
100
VEH
MDL
M
HDL Size
0.0933
nm
9.0
9.5
10.0
10.5
11.0
11.5
VEH
MDL
N
TRLTG
0.0506
mg/dL
0
10
20
30
40
50
VEH
MDL
O
TRLC
0.0752
mg/dL
0
5
10
15
VEH
MDL

**Figure S.4**

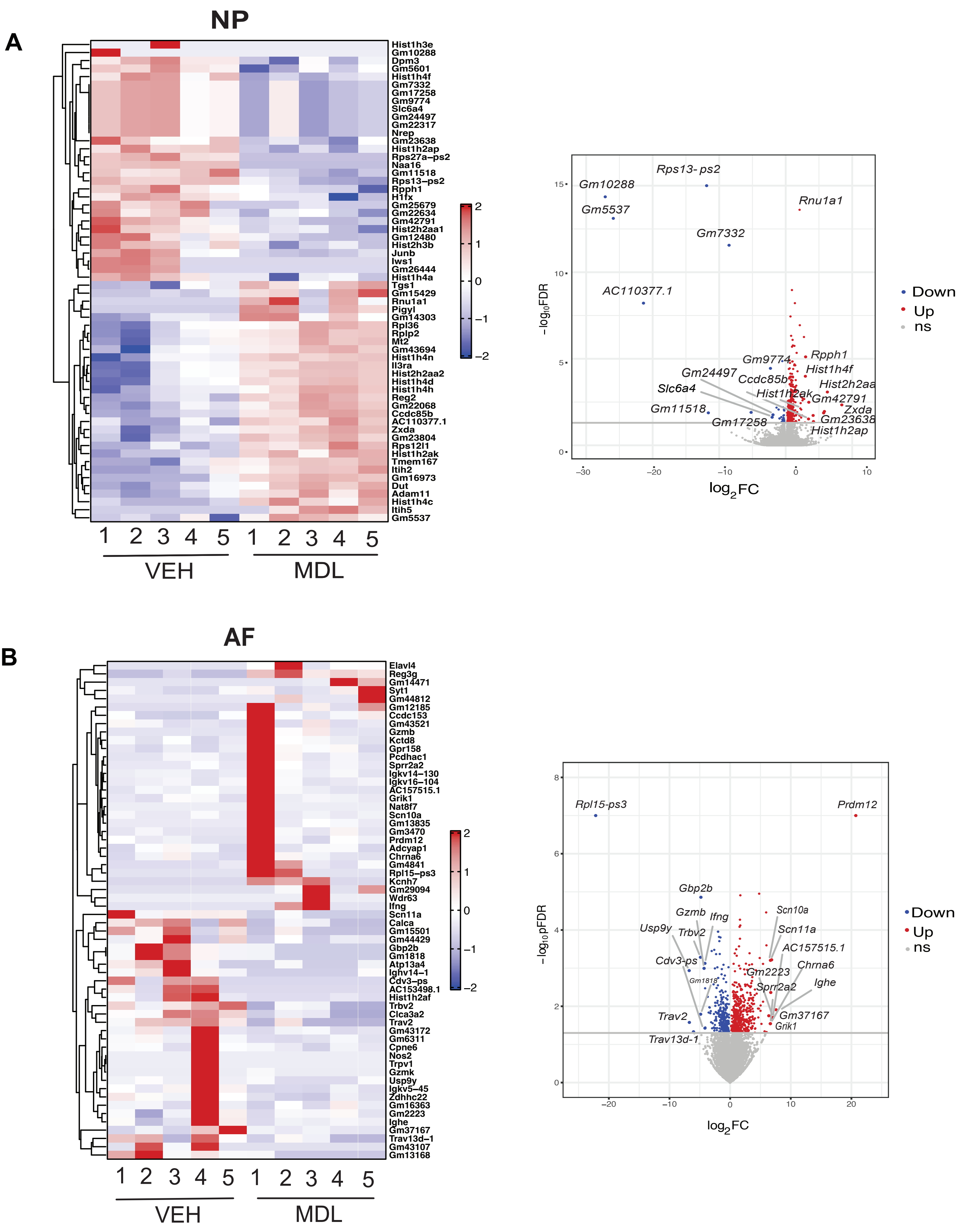
A
NP
Hist1h3e
Gm10288
Dpm3
Gm5601
Hist1h4f
Gm7332
Gm17258
Gm9774
Slc6a4
Gm24497
Gm22317
Nrep
Gm23638
Hist1h2ap
Rps27a-ps2
Naa16
Gm11518
Rps13-ps2
Rpph1
H1fx
Gm25679
Gm22634
Gm42791
Hist2h2aa1
Gm12480
Hist2h3b
Junb
Iws1
Gm26444
Hist1h4a
Tgs1
Gm15429
Rnu1a1
Pigyl
Gm14303
Rpl36
Rplp2
Mt2
Gm43694
Hist1h4n
Il3ra
Hist2h2aa2
Hist1h4d
Hist1h4h
Reg2
Gm22068
Ccdc85b
AC110377.1
Zxda
Gm23804
Rps12l1
Hist1h2ak
Tmem167
Itih2
Gm16973
Dut
Adam11
Hist1h4c
Itih5
Gm5537
2
1
0
−1
−2
1 2 3 4 5 1 2 3 4 5
VEH
MDL
Rps13- ps2
Gm10288
Gm5537
Rnu1a1
Gm7332
AC110377.1
Gm9774
Rpph1
Hist1h4f
Gm24497
Ccdc85b
Hist2h2aa
Slc6a4
Hist1h2ak
Gm42791
Gm11518
Zxda
Gm17258
Gm23638
Hist1h2ap
15
10
5
0
−30
−20
−10
0
10
−log10FDR
log2FC
Down
Up
ns
B
AF
Elavl4
Reg3g
Gm14471
Syt1
Gm44812
Gm12185
Ccdc153
Gm43521
Gzmb
Kctd8
Gpr158
Pcdhac1
Sprr2a2
Igkv14−130
Igkv16−104
AC157515.1
Grik1
Nat8f7
Scn10a
Gm13835
Gm3470
Prdm12
Adcyap1
Chrna6
Gm4841
Rpl15−ps3
Kcnh7
Gm29094
Wdr63
Ifng
Scn11a
Calca
Gm15501
Gm44429
Gbp2b
Gm1818
Atp13a4
Ighv14−1
Cdv3−ps
AC153498.1
Hist1h2af
Trbv2
Clca3a2
Trav2
Gm43172
Gm6311
Cpne6
Nos2
Trpv1
Gzmk
Usp9y
Igkv5−45
Zdhhc22
Gm16363
Gm2223
Ighe
Gm37167
Trav13d−1
Gm43107
Gm13168
2
1
0
−1
−2
1 2 3 4 5 1 2 3 4 5
VEH
MDL
Rpl15-ps3
Prdm12
Gbp2b
Gzmb
Ifng
Scn10a
Usp9y
Trbv2
Scn11a
AC157515.1
Cdv3-ps
Chrna6
Gm2223
Gm1818
Sprr2a2
Ighe
Trav2
Gm37167
Grik1
Trav13d-1
8
6
4
2
0
−20
−10
0
10
20
−log10pFDR
log2FC
Down
Up
ns

Figure S.5

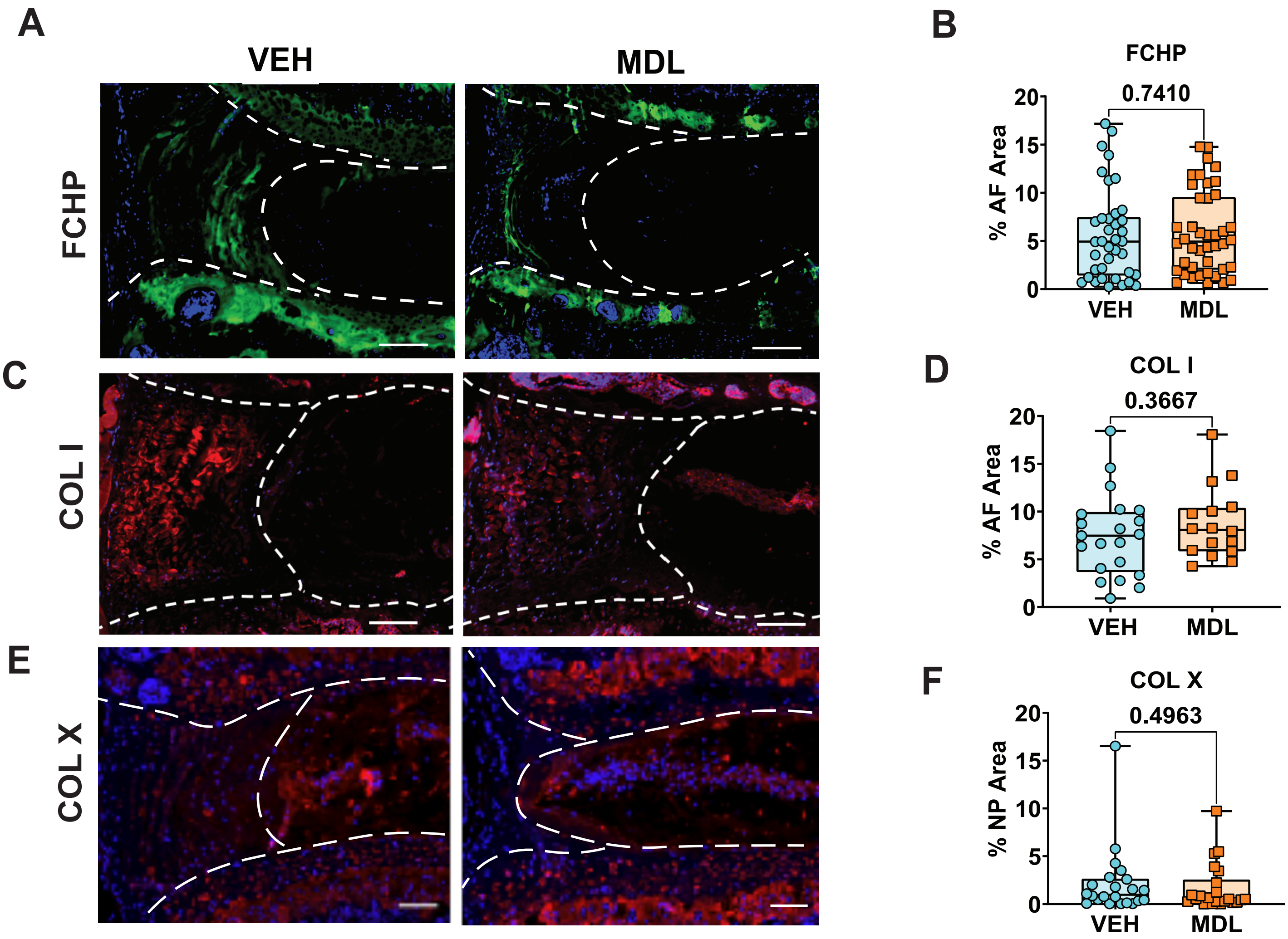
A
VEH
MDL
FCHP
B
FCHP
0.7410
% AF Area
20
15
10
5
0
VEH
MDL
C
COL I
D
COL I
0.3667
% AF Area
20
15
10
5
0
VEH
MDL
E
COL X
F
COL X
0.4963
% NP Area
20
15
10
5
0
VEH
MDL

**Figure S.6**

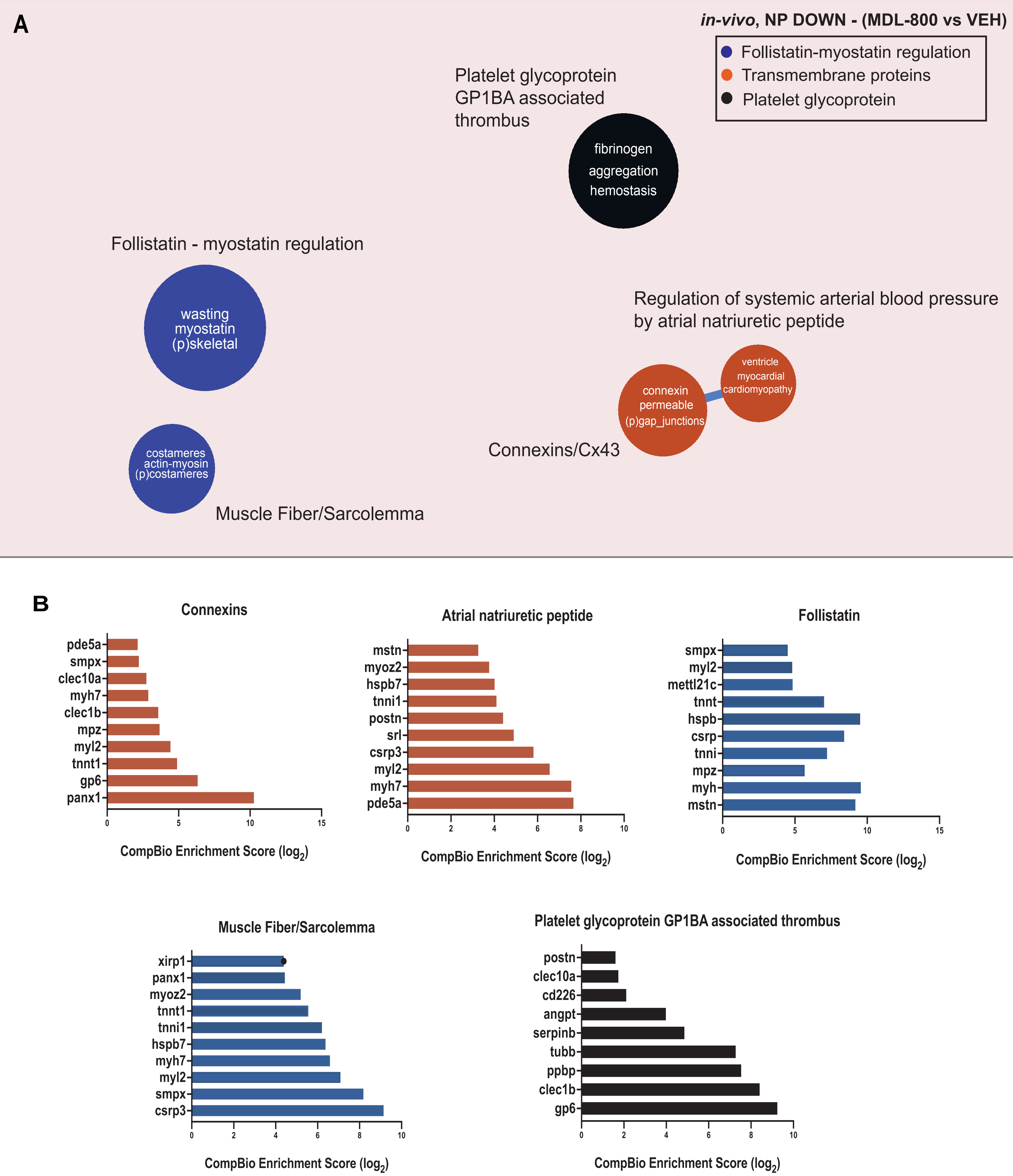

A
in-vivo, NP DOWN - (MDL-800 vs VEH)
Follistatin-myostatin regulation
Transmembrane proteins
Platelet glycoprotein
Platelet glycoprotein GP1BA associated thrombus
fibrinogen
aggregation
hemostasis
Follistatin - myostatin regulation
wasting
myostatin
(p)skeletal
Regulation of systemic arterial blood pressure by atrial natriuretic peptide
ventricle
myocardial
cardiomyopathy
connexin
permeable
(p)gap_junctions
Connexins/Cx43
costameres
actin-myosin
(p)costameres
Muscle Fiber/Sarcolemma
B
Connexins
pde5a
smpx
clec10a
myh7
clec1b
mpz
myl2
tnnt1
gp6
panx1
0
5
10
15
CompBio Enrichment Score (log2)
Atrial natriuretic peptide
mstn
myoz2
hspb7
tnni1
postn
srl
csrp3
myl2
myh7
pde5a
0
2
4
6
8
10
CompBio Enrichment Score (log2)
Follistatin
smpx
myl2
mettl21c
tnnt
hspb
csrp
tnni
mpz
myh
mstn
0
5
10
15
CompBio Enrichment Score (log2)
Muscle Fiber/Sarcolemma
xirp1
panx1
myoz2
tnnt1
tnni1
hspb7
myh7
myl2
smpx
csrp3
0
2
4
6
8
10
CompBio Enrichment Score (log2)
Platelet glycoprotein GP1BA associated thrombus
postn
clec10a
cd226
angpt
serpinb
tubb
ppbp
clec1b
gp6
0
2
4
6
8
10
CompBio Enrichment Score (log2)

Figure S.7

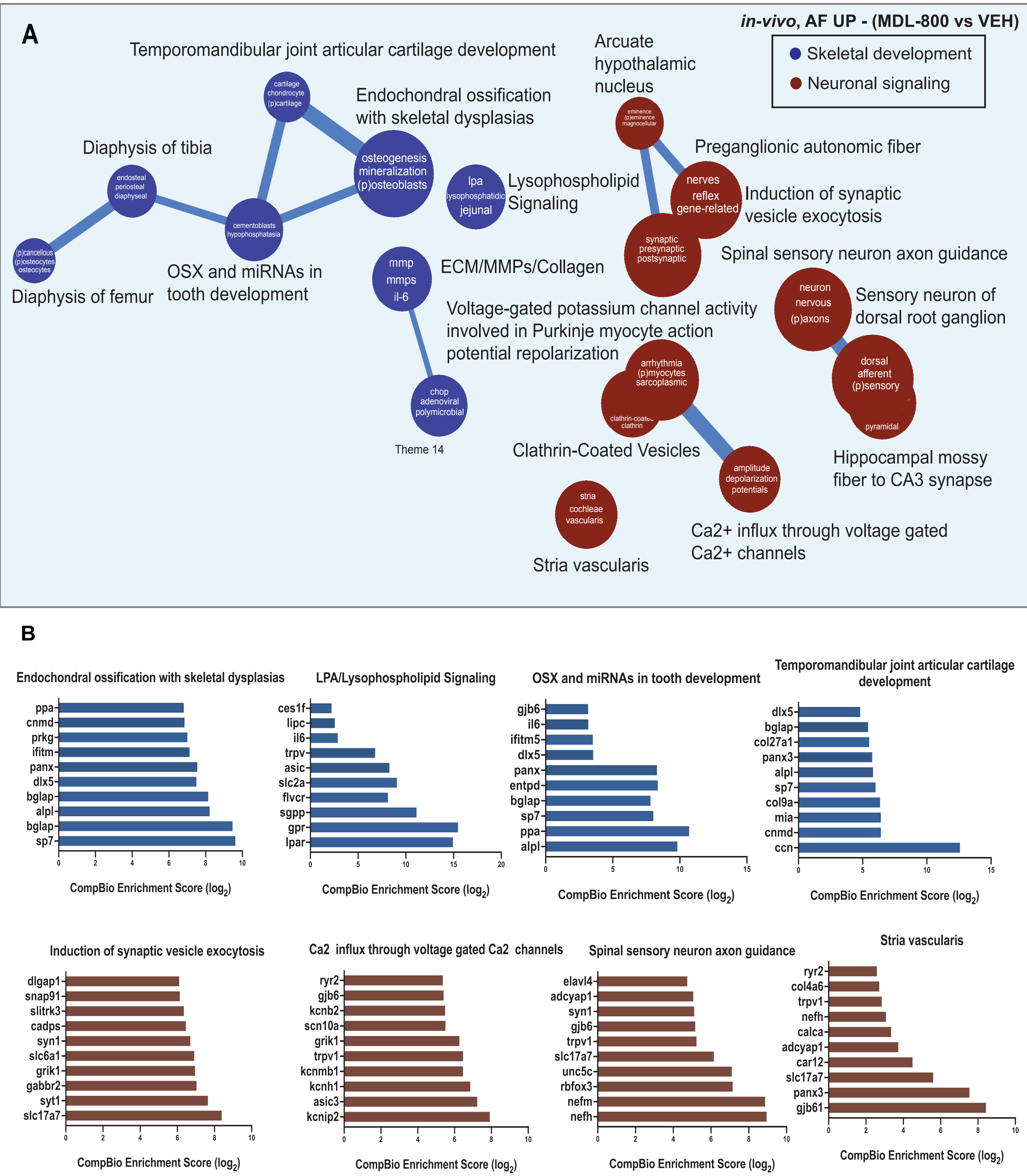
A
in-vivo, AF UP - (MDL-800 vs VEH)
Skeletal development
Neuronal signaling
Temporomandibular joint articular cartilage development
cartilage chondrocyte (p)cartilage
Endochondral ossification with skeletal dysplasias
osteogenesis mineralization (p)osteoblasts
Diaphysis of tibia
endosteal periosteal diaphyseal
cementoblasts hypophosphatasia
(p)cancellous (p)osteocytes osteocytes
Diaphysis of femur
OSX and miRNAs in tooth development
lpa lysophosphatidic jejunal
Lysophospholipid Signaling
mmp mmps il-6
ECM/MMPs/Collagen
chop adenoviral polymicrobial
Theme 14
Arcuate hypothalamic nucleus
eminence (p)eminence magnocellular
Preganglionic autonomic fiber
nerves reflex gene-related
Induction of synaptic vesicle exocytosis
synaptic presynaptic postsynaptic
Spinal sensory neuron axon guidance
neuron nervous (p)axons
Sensory neuron of dorsal root ganglion
Voltage-gated potassium channel activity involved in Purkinje myocyte action potential repolarization
arrhythmia (p)myocytes sarcoplasmic
clathrin-coated clathrin
Clathrin-Coated Vesicles
dorsal afferent (p)sensory
pyramidal
Hippocampal mossy fiber to CA3 synapse
amplitude depolarization potentials
stria cochleae vascularis
Ca2+ influx through voltage gated Ca2+ channels
Stria vascularis
B
Endochondral ossification with skeletal dysplasias
ppa
cnmd
prkg
ifitm
panx
dlx5
bglap
alpl
bglap
sp7
CompBio Enrichment Score (log2)
LPA/Lysophospholipid Signaling
ces1f
lipc
il6
trpv
asic
slc2a
flvcr
sgpp
gpr
lpar
CompBio Enrichment Score (log2)
OSX and miRNAs in tooth development
gjb6
il6
ifitm5
dlx5
panx
entpd
bglap
sp7
ppa
alpl
CompBio Enrichment Score (log2)
Temporomandibular joint articular cartilage development
dlx5
bglap
col27a1
panx3
alpl
sp7
col9a
mia
cnmd
ccn
CompBio Enrichment Score (log2)
Induction of synaptic vesicle exocytosis
dlgap1
snap91
slitrk3
cadps
syn1
slc6a1
grik1
gabbr2
syt1
slc17a7
CompBio Enrichment Score (log2)
Ca2 influx through voltage gated Ca2 channels
ryr2
gjb6
kcnb2
scn10a
grik1
trpv1
kcnmb1
kcnh1
asic3
kcnip2
CompBio Enrichment Score (log2)
Spinal sensory neuron axon guidance
elavl4
adcyap1
syn1
gjb6
trpv1
slc17a7
unc5c
rbfox3
nefm
nefh
CompBio Enrichment Score (log2)
Stria vascularis
ryr2
col4a6
trpv1
nefh
calca
adcyap1
car12
slc17a7
panx3
gjb61
CompBio Enrichment Score (log2)

#### Figure S.8

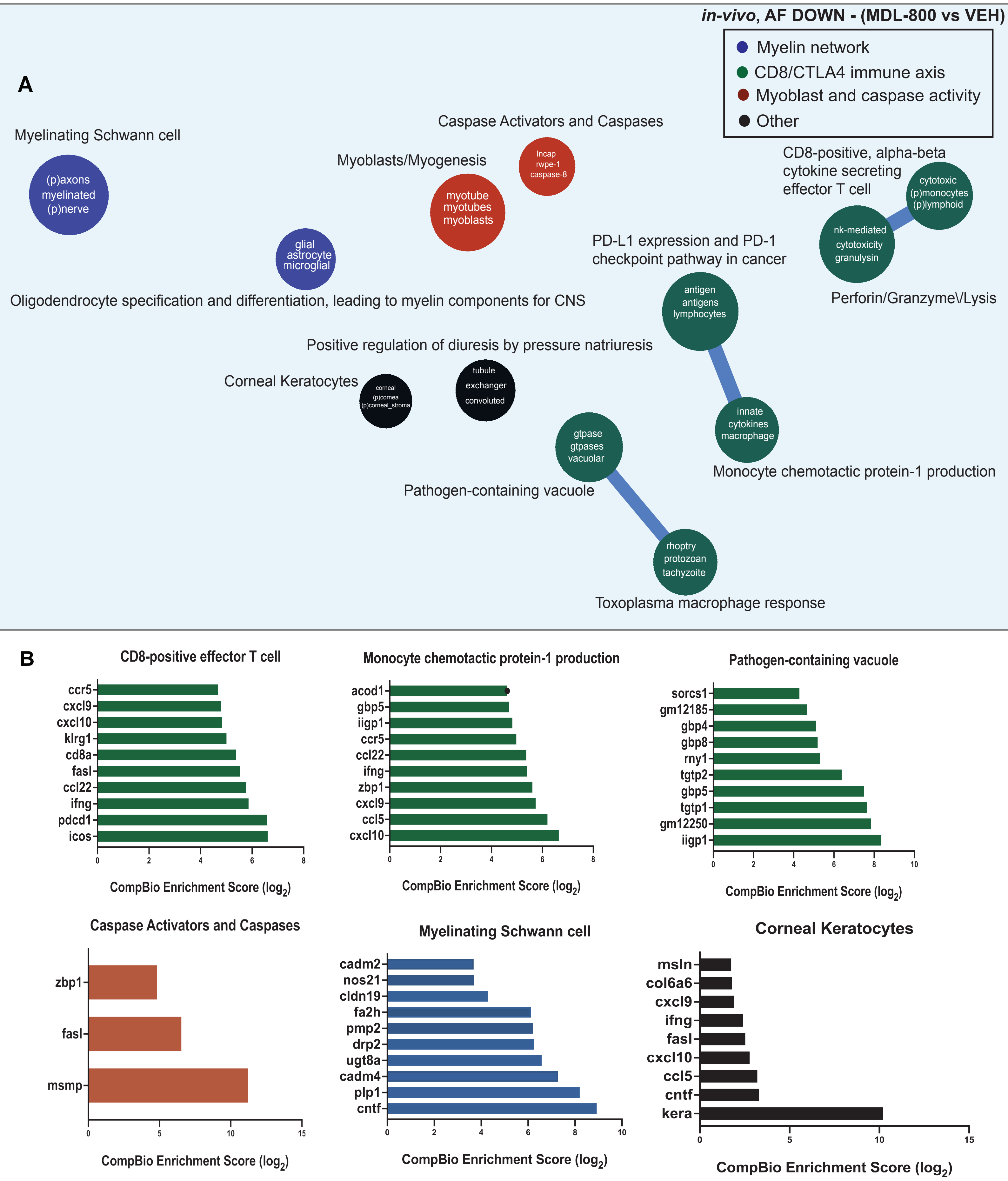
in-vivo, AF DOWN - (MDL-800 vs VEH)
Myelin network
CD8/CTLA4 immune axis
Myoblast and caspase activity
Other
A
Myelinating Schwann cell
(p)axons
myelinated
(p)nerve
Caspase Activators and Caspases
lncap
rwpe-1
caspase-8
Myoblasts/Myogenesis
myotube
myotubes
myoblasts
CD8-positive, alpha-beta
cytokine secreting
effector T cell
cytotoxic
(p)monocytes
(p)lymphoid
nk-mediated
cytotoxicity
granulysin
Perforin/GranzymeV/Lysis
glial
astrocyte
microglial
Oligodendrocyte specification and differentiation, leading to myelin components for CNS
PD-L1 expression and PD-1
checkpoint pathway in cancer
antigen
antigens
lymphocytes
Positive regulation of diuresis by pressure natriuresis
Corneal Keratocytes
corneal
(p)cornea
(p)corneal_stroma
tubule
exchanger
convoluted
innate
cytokines
macrophage
gtpase
gtpases
vacuolar
Monocyte chemotactic protein-1 production
Pathogen-containing vacuole
rhoptry
protozoan
tachyzoite
Toxoplasma macrophage response
B
CD8-positive effector T cell
ccr5
cxcl9
cxcl10
klrg1
cd8a
fasl
ccl22
ifng
pdcd1
icos
0
2
4
6
8
CompBio Enrichment Score (log2)
Monocyte chemotactic protein-1 production
acod1
gbp5
iigp1
ccr5
ccl22
ifng
zbp1
cxcl9
ccl5
cxcl10
0
2
4
6
8
CompBio Enrichment Score (log2)
Pathogen-containing vacuole
sorcs1
gm12185
gbp4
gbp8
rny1
tgtp2
gbp5
tgtp1
gm12250
iigp1
0
2
4
6
8
10
CompBio Enrichment Score (log2)
Caspase Activators and Caspases
zbp1
fasl
msmp
0
5
10
15
CompBio Enrichment Score (log2)
Myelinating Schwann cell
cadm2
nos21
cldn19
fa2h
pmp2
drp2
ugt8a
cadm4
plp1
cntf
0
2
4
6
8
10
CompBio Enrichment Score (log2)
Corneal Keratocytes
msln
col6a6
cxcl9
ifng
fasl
cxcl10
ccl5
cntf
kera
0
5
10
15
CompBio Enrichment Score (log2)

**Figure S.9**

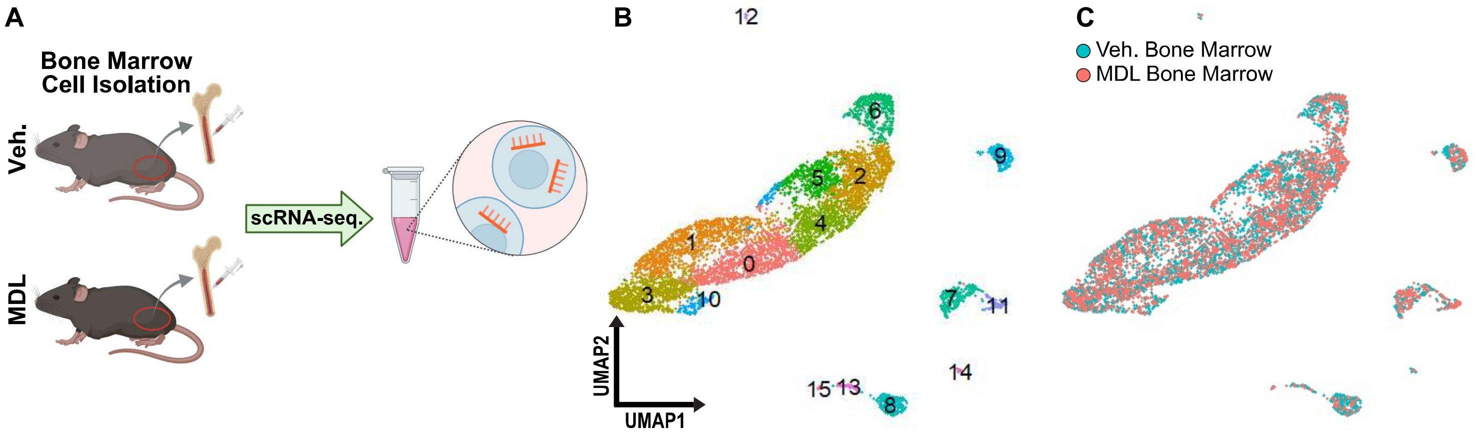


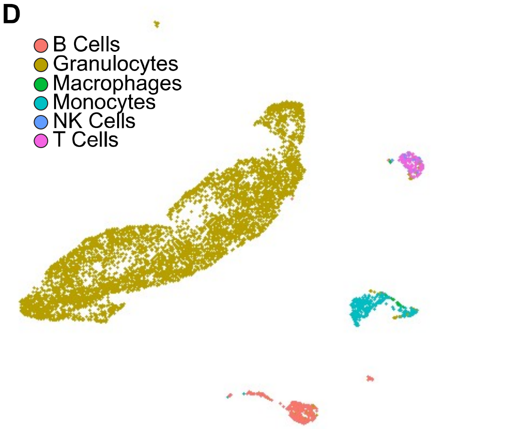


E

| Annotation | Cell Count Veh. | Cell Count MDL | Percent Veh. | Percent MDL | OR | p-value |
|---|---|---|---|---|---|---|
| B Cells | 279 | 76 | 7.69% | 2.58% | 0.32 | 5.6e-21 |
| Granulocytes | 3026 | 2616 | 83.45% | 88.89% | 1.59 | 2.6e-10 |
| Macrophages | 4 | 12 | 0.14% | 0.41% | 2.96 | 0.048 |
| Monocytes | 193 | 175 | 5.32% | 5.95% | 1.12 | 0.28 |
| NK Cells | 23 | 10 | 0.63% | 0.34% | 0.53 | 0.11 |
| T Cells | 100 | 54 | 2.76% | 1.83% | 0.66 | 0.014 |
| Total | 3626 | 2943 | 100% | 100% | | |

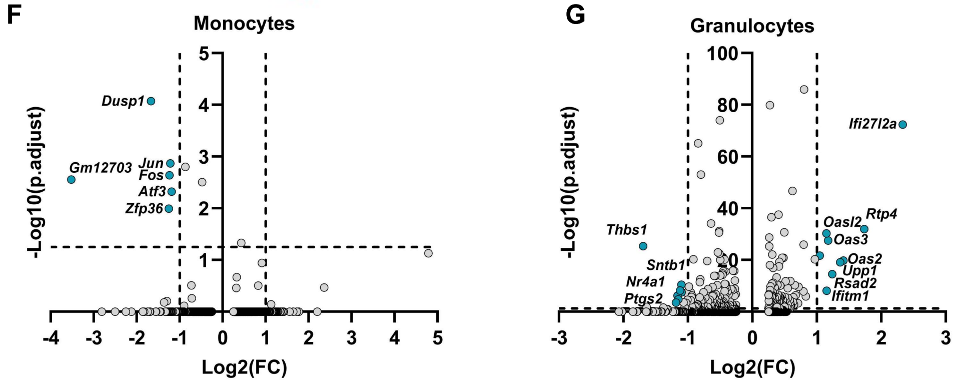

**Figure S.10**

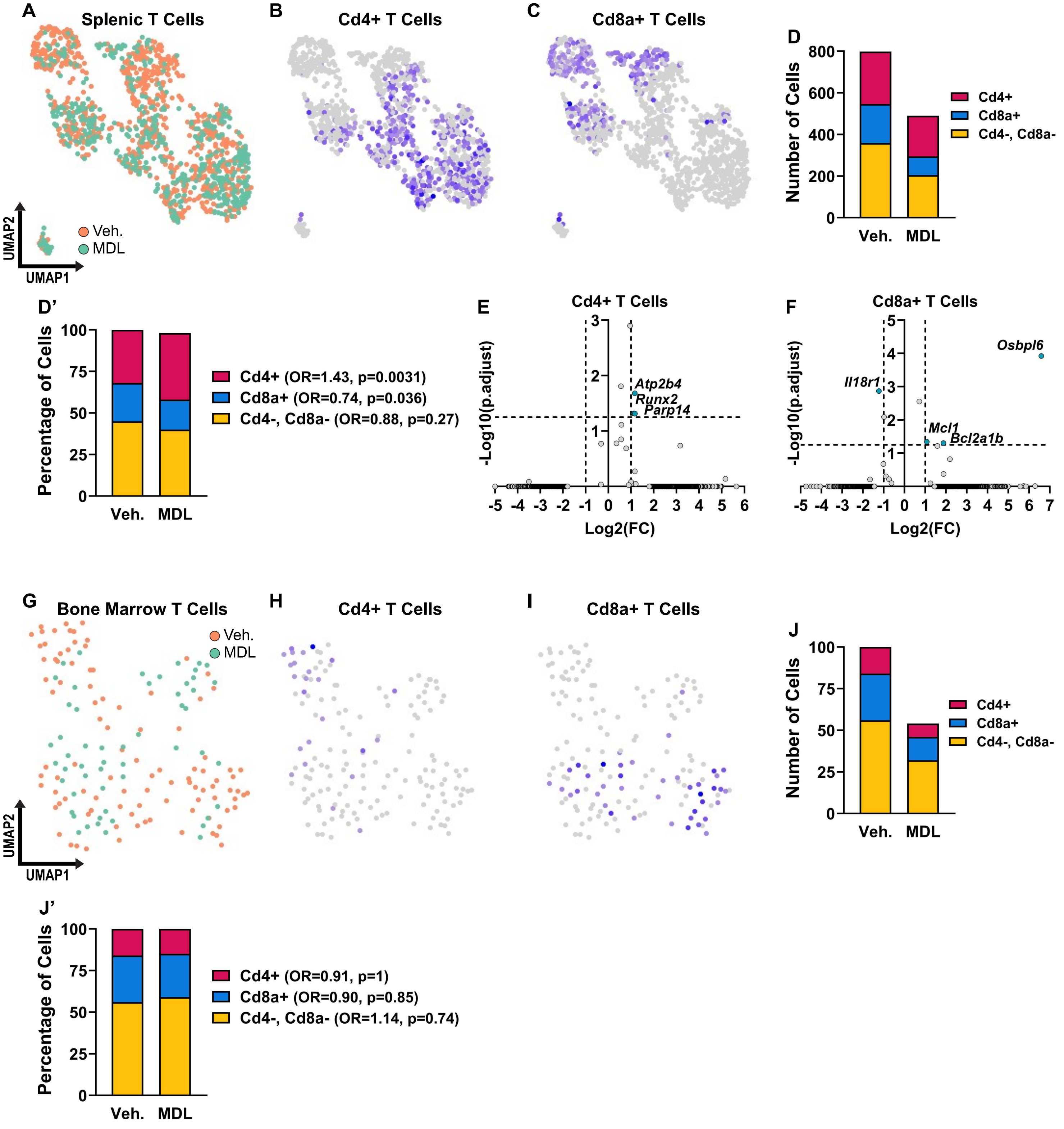
A
Splenic T Cells
Veh.
MDL
UMAP2
UMAP1
B
Cd4+ T Cells
C
Cd8a+ T Cells
D
Number of Cells
800
600
400
200
0
Veh.
MDL
Cd4+
Cd8a+
Cd4-, Cd8a-
D'
Percentage of Cells
100
75
50
25
0
Veh.
MDL
Cd4+ (OR=1.43, p=0.0031)
Cd8a+ (OR=0.74, p=0.036)
Cd4-, Cd8a- (OR=0.88, p=0.27)
E
Cd4+ T Cells
-Log10(p.adjust)
Atp2b4
Runx2
Parp14
Log2(FC)
-5 -4 -3 -2 -1 0 1 2 3 4 5 6
F
Cd8a+ T Cells
-Log10(p.adjust)
Osbpl6
Il18r1
Mcl1
Bcl2a1b
Log2(FC)
-5 -4 -3 -2 -1 0 1 2 3 4 5 6 7
G
Bone Marrow T Cells
Veh.
MDL
UMAP2
UMAP1
H
Cd4+ T Cells
I
Cd8a+ T Cells
J
Number of Cells
100
75
50
25
0
Veh.
MDL
Cd4+
Cd8a+
Cd4-, Cd8a-
J'
Percentage of Cells
100
75
50
25
0
Veh.
MDL
Cd4+ (OR=0.91, p=1)
Cd8a+ (OR=0.90, p=0.85)
Cd4-, Cd8a- (OR=1.14, p=0.74)

Figure S.11

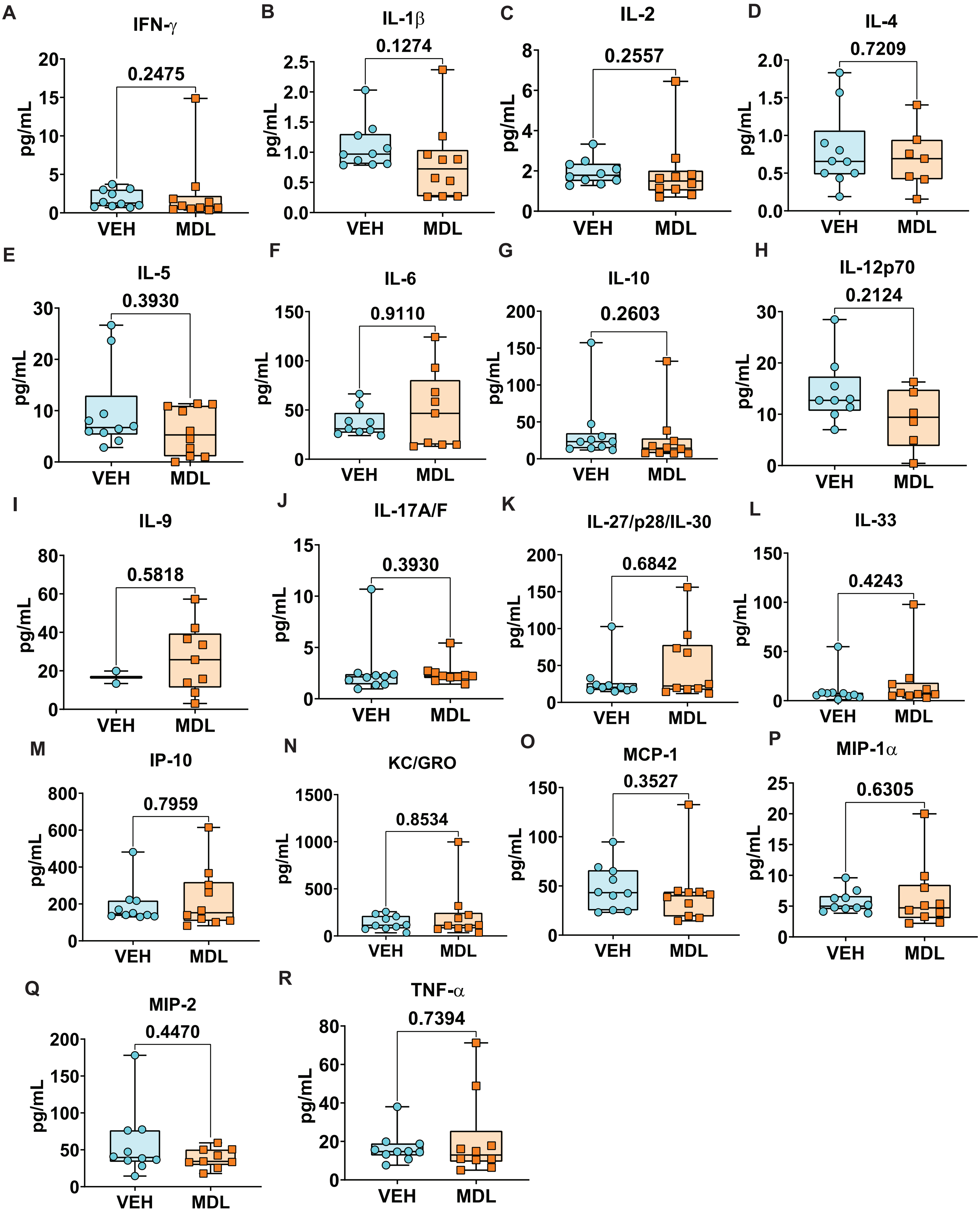
A
IFN-γ
0.2475
B
IL-1β
0.1274
C
IL-2
0.2557
D
IL-4
0.7209
E
IL-5
0.3930
F
IL-6
0.9110
G
IL-10
0.2603
H
IL-12p70
0.2124
I
IL-9
0.5818
J
IL-17A/F
0.3930
K
IL-27/p28/IL-30
0.6842
L
IL-33
0.4243
M
IP-10
0.7959
N
KC/GRO
0.8534
O
MCP-1
0.3527
P
MIP-1α
0.6305
Q
MIP-2
0.4470
R
TNF-α
0.7394
pg/mL
VEH
MDL

Figure S.12

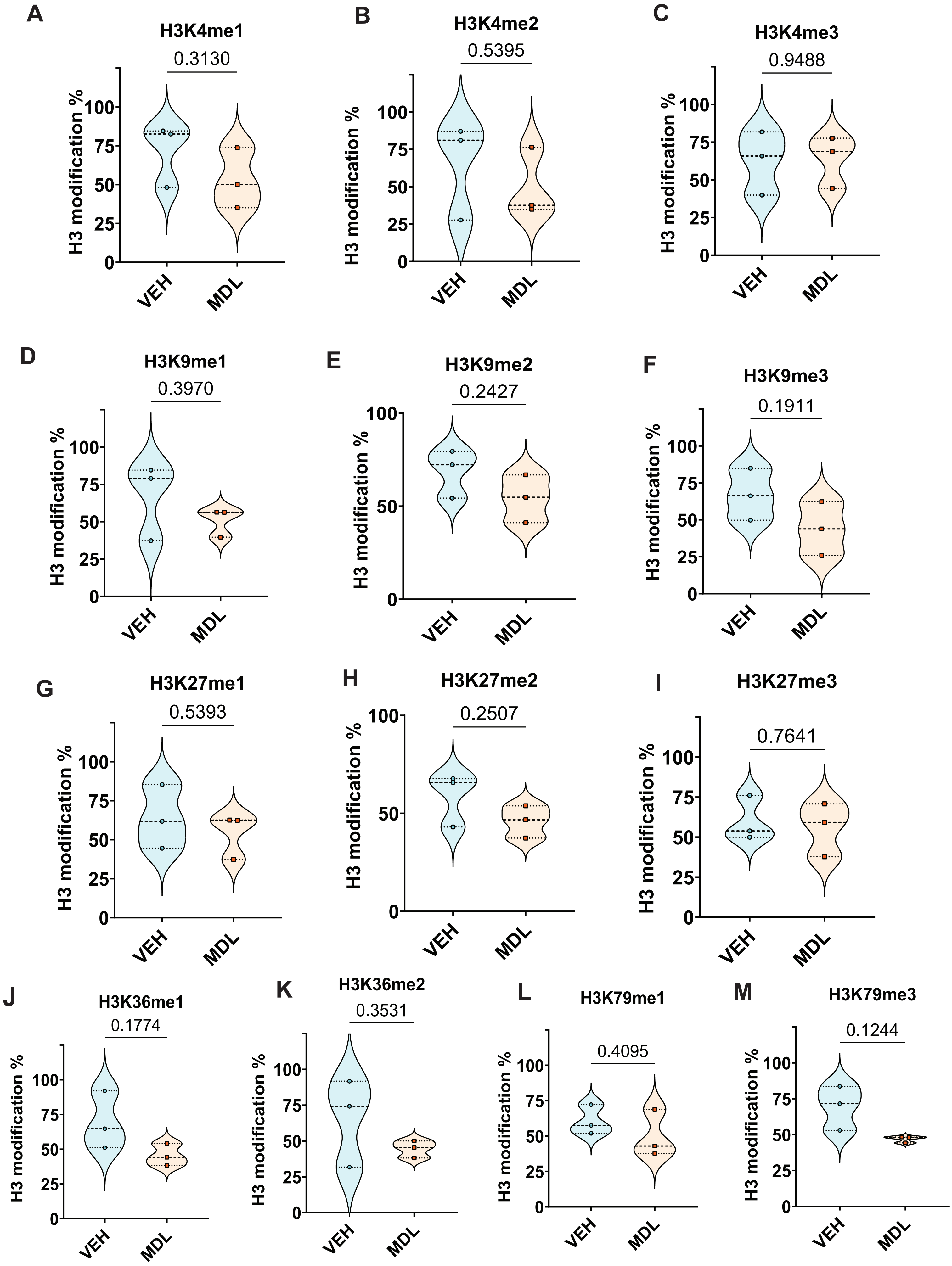
A
H3K4me1
0.3130
B
H3K4me2
0.5395
C
H3K4me3
0.9488
D
H3K9me1
0.3970
E
H3K9me2
0.2427
F
H3K9me3
0.1911
G
H3K27me1
0.5393
H
H3K27me2
0.2507
I
H3K27me3
0.7641
J
H3K36me1
0.1774
K
H3K36me2
0.3531
L
H3K79me1
0.4095
M
H3K79me3
0.1244
H3 modification %
VEH
MDL

**Figure S.13**

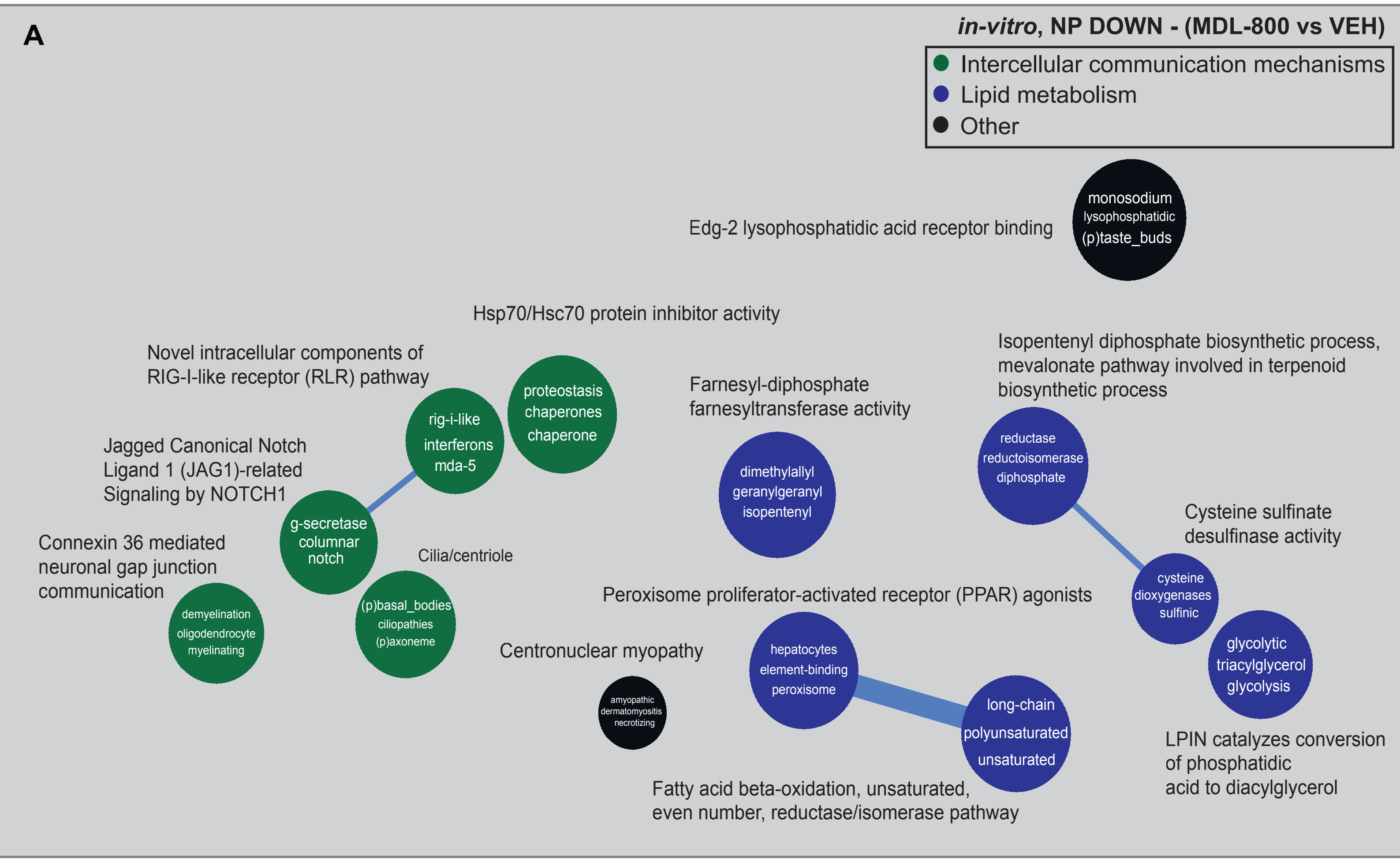
A
in-vitro, NP DOWN - (MDL-800 vs VEH)
Intercellular communication mechanisms
Lipid metabolism
Other
Edg-2 lysophosphatidic acid receptor binding
monosodium lysophosphatidic (p)taste_buds
Hsp70/Hsc70 protein inhibitor activity
proteostasis chaperones chaperone
Novel intracellular components of RIG-I-like receptor (RLR) pathway
rig-i-like interferons mda-5
Jagged Canonical Notch Ligand 1 (JAG1)-related Signaling by NOTCH1
g-secretase columnar notch
Connexin 36 mediated neuronal gap junction communication
demyelination oligodendrocyte myelinating
Cilia/centriole
(p)basal_bodies ciliopathies (p)axoneme
Farnesyl-diphosphate farnesyltransferase activity
dimethylallyl geranylgeranyl isopentenyl
Isopentenyl diphosphate biosynthetic process, mevalonate pathway involved in terpenoid biosynthetic process
reductase reductoisomerase diphosphate
Cysteine sulfinate desulfinase activity
cysteine dioxygenases sulfinic
Peroxisome proliferator-activated receptor (PPAR) agonists
hepatocytes element-binding peroxisome
Centronuclear myopathy
amyopathic dermatomyositis necrotizing
long-chain polyunsaturated unsaturated
glycolytic triacylglycerol glycolysis
LPIN catalyzes conversion of phosphatidic acid to diacylglycerol
Fatty acid beta-oxidation, unsaturated, even number, reductase/isomerase pathway

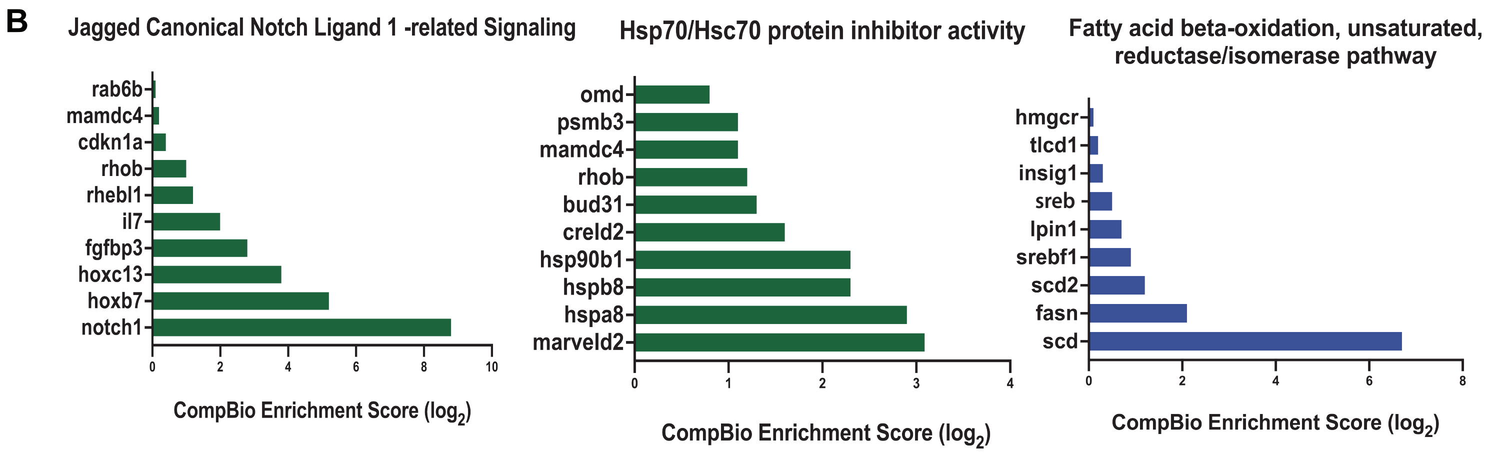
B
Jagged Canonical Notch Ligand 1 -related Signaling
rab6b
mamdc4
cdkn1a
rhob
rhebl1
il7
fgfbp3
hoxc13
hoxb7
notch1
0 2 4 6 8 10
CompBio Enrichment Score (log2)
Hsp70/Hsc70 protein inhibitor activity
omd
psmb3
mamdc4
rhob
bud31
creld2
hsp90b1
hspb8
hspa8
marveld2
0 1 2 3 4
CompBio Enrichment Score (log2)
Fatty acid beta-oxidation, unsaturated, reductase/isomerase pathway
hmgcr
tlcd1
insig1
sreb
lpin1
srebf1
scd2
fasn
scd
0 2 4 6 8
CompBio Enrichment Score (log2)

### Figure S.14

**A**

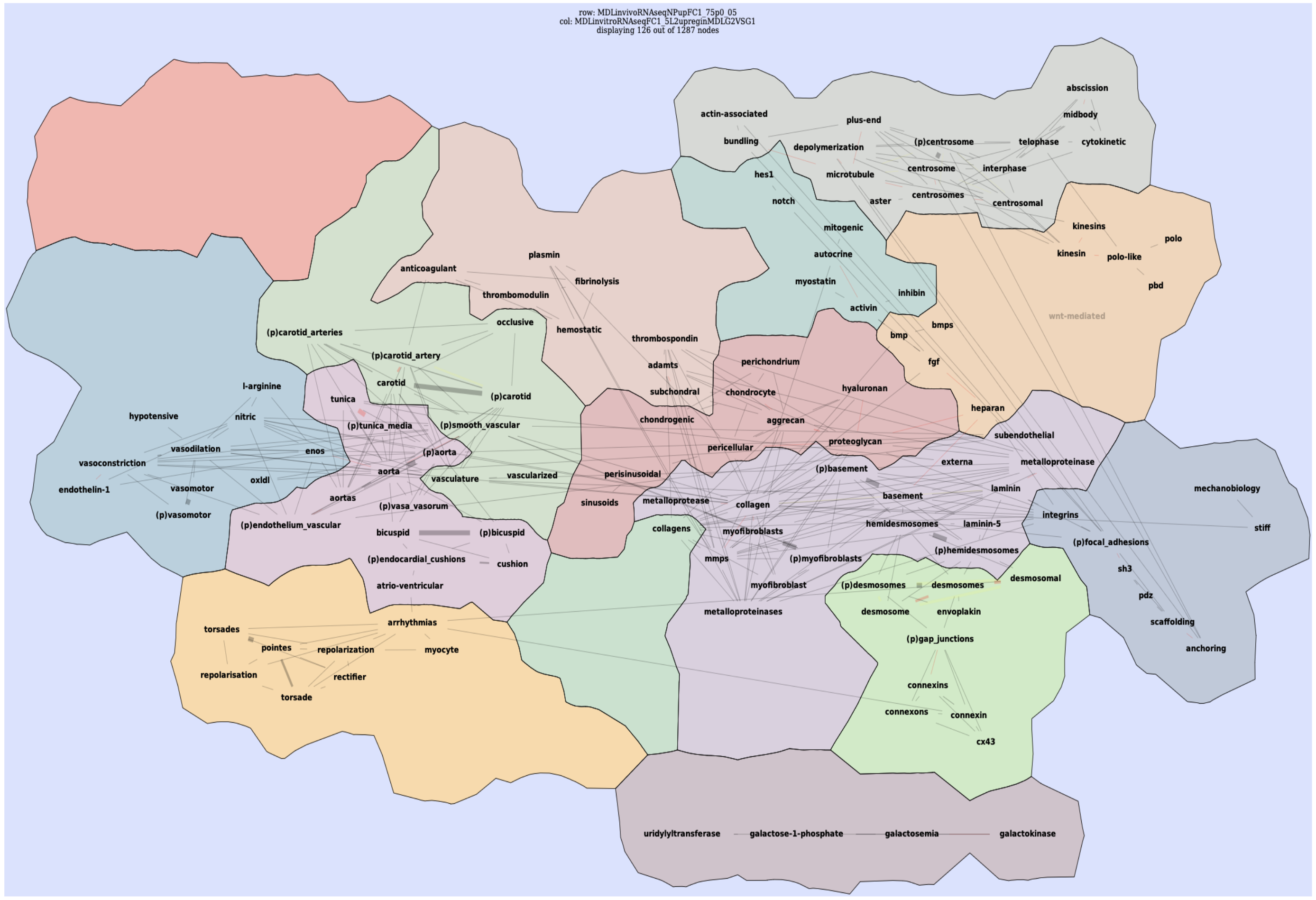


**B**

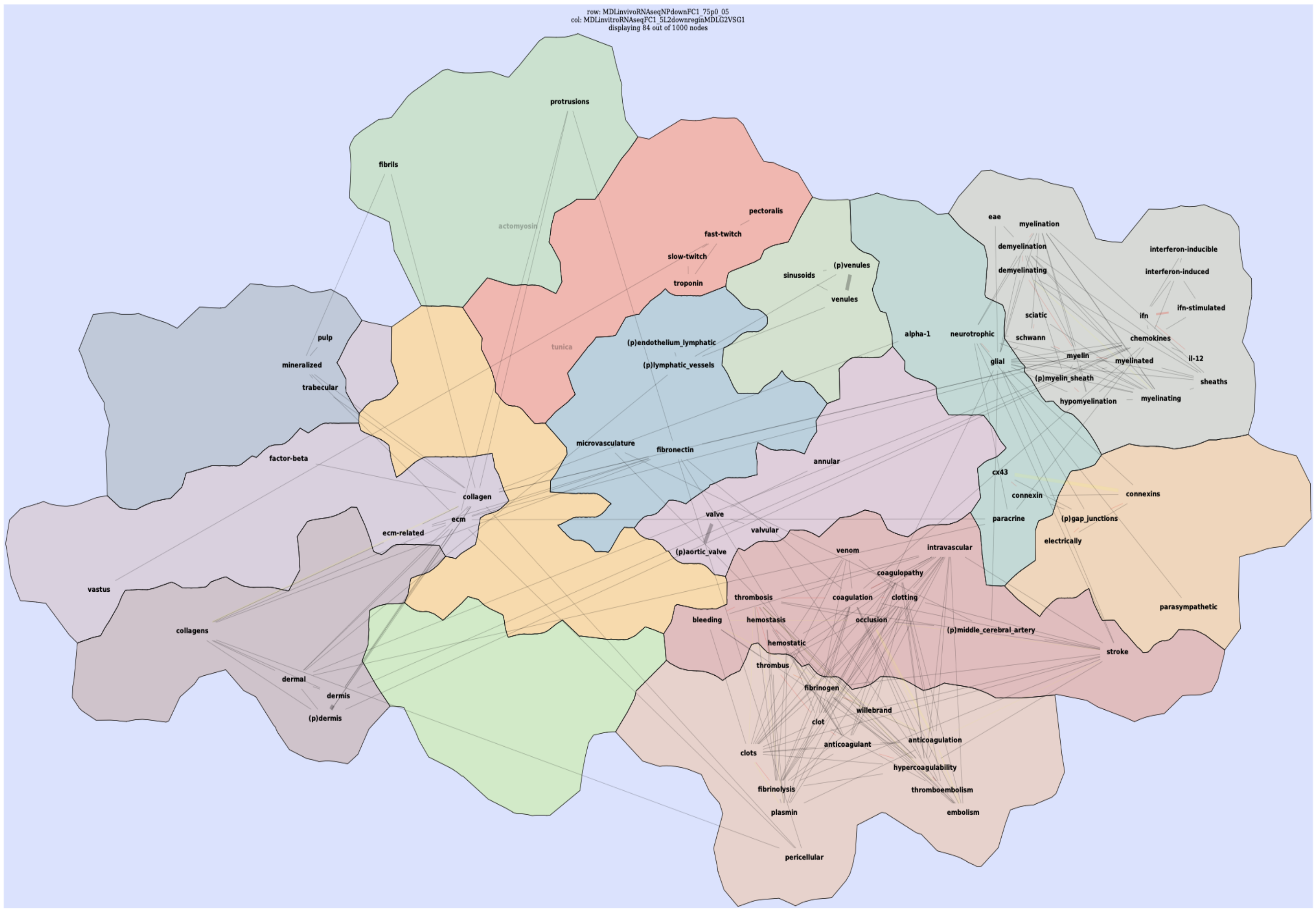

Figure S.15

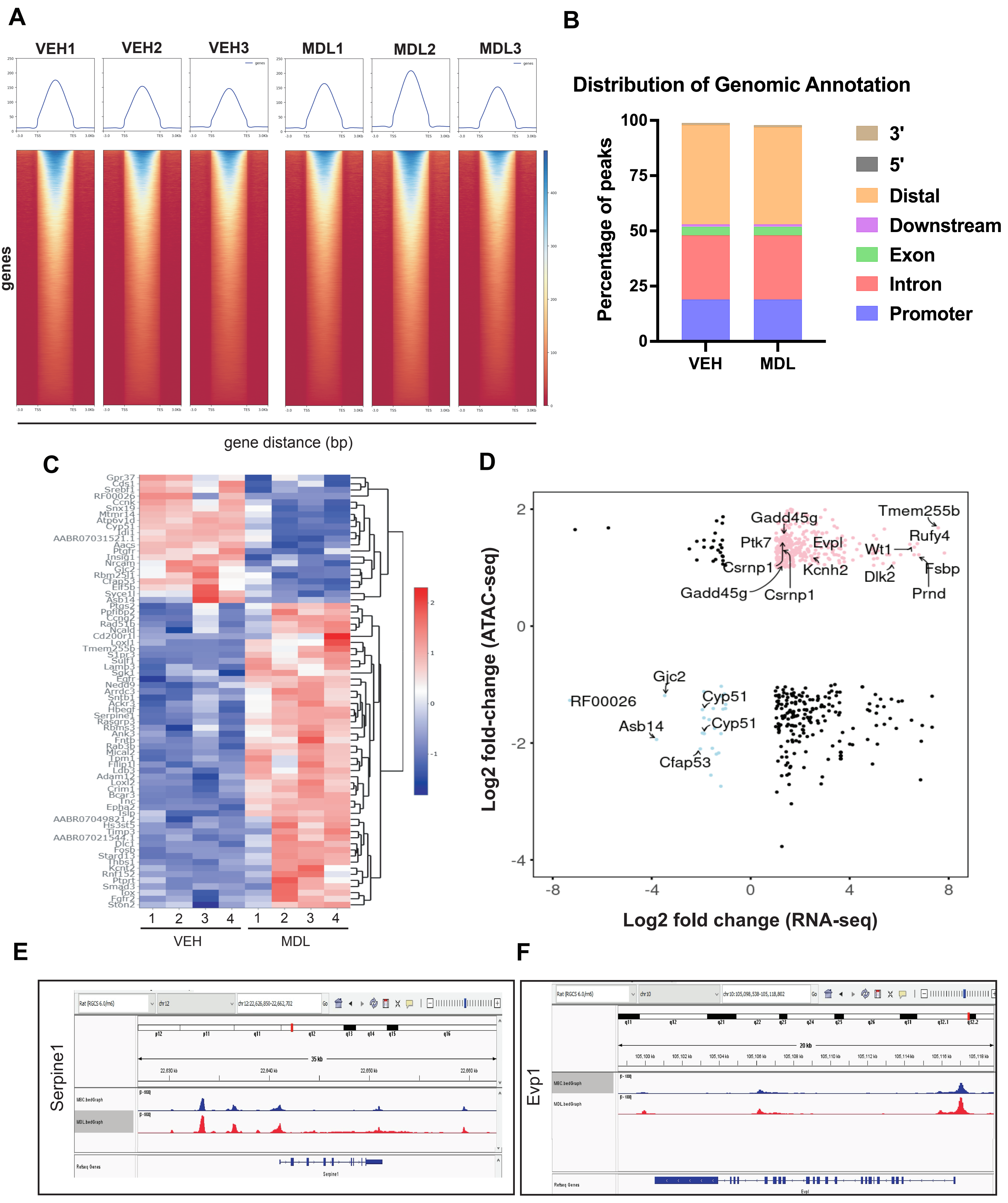
A
VEH1
VEH2
VEH3
MDL1
MDL2
MDL3
genes
gene distance (bp)
B
Distribution of Genomic Annotation
Percentage of peaks
VEH
MDL
3'
5'
Distal
Downstream
Exon
Intron
Promoter
C
VEH
MDL
D
Log2 fold-change (ATAC-seq)
Log2 fold change (RNA-seq)
Gadd45g
Tmem255b
Ptk7
Evpl
Rufy4
Wt1
Csrnp1
Kcnh2
Dlk2
Fsbp
Prnd
Gjc2
RF00026
Cyp51
Asb14
Cfap53
E
Serpine1
F
Evp1

**Figure S.16**

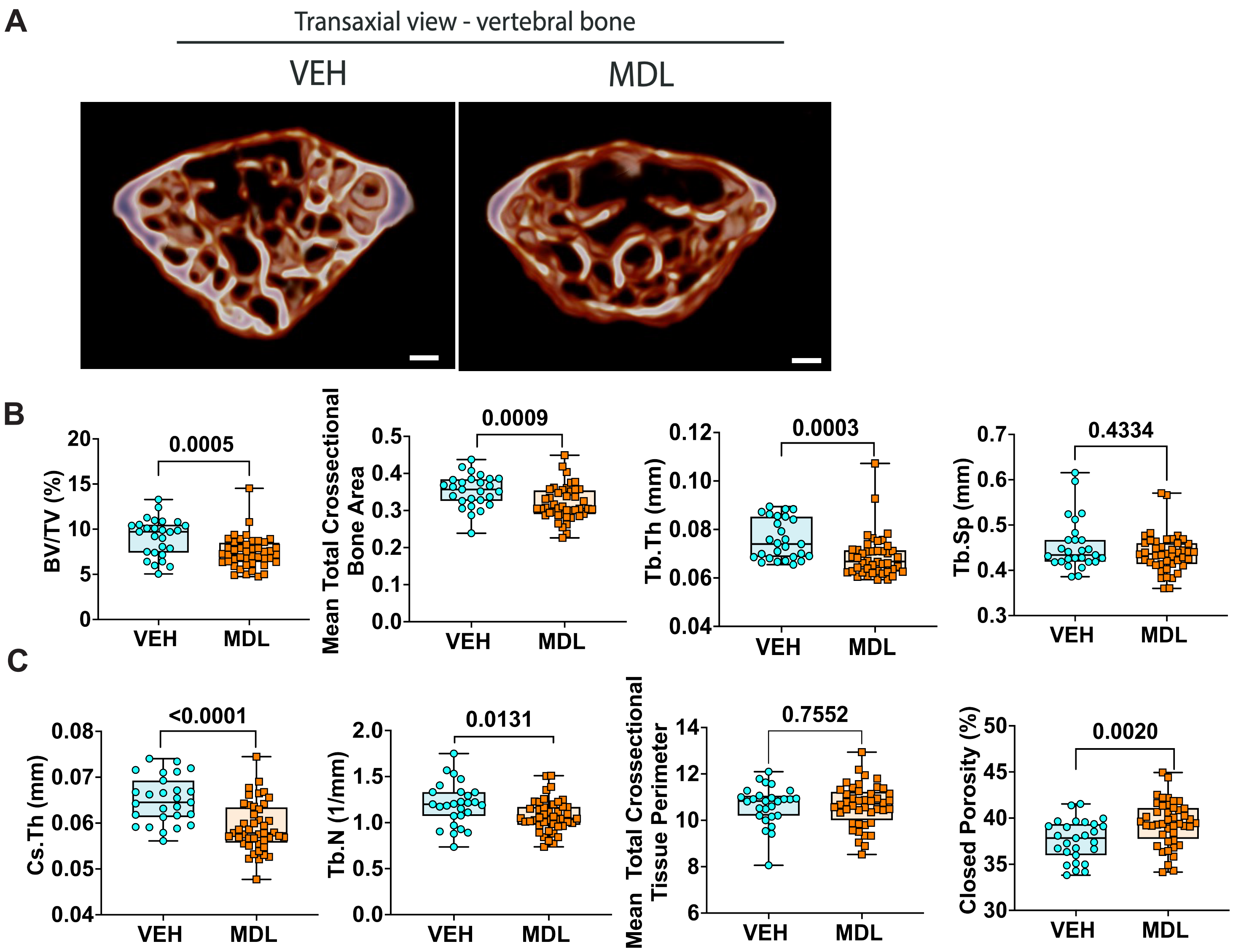
A
Transaxial view - vertebral bone
VEH
MDL
B
BV/TV (%)
0.0005
Mean Total Crossectional Bone Area
0.0009
Tb.Th (mm)
0.0003
Tb.Sp (mm)
0.4334
C
Cs.Th (mm)
<0.0001
Tb.N (1/mm)
0.0131
Mean Total Crossectional Tissue Perimeter
0.7552
Closed Porosity (%)
0.0020
VEH
MDL

Figure S.17

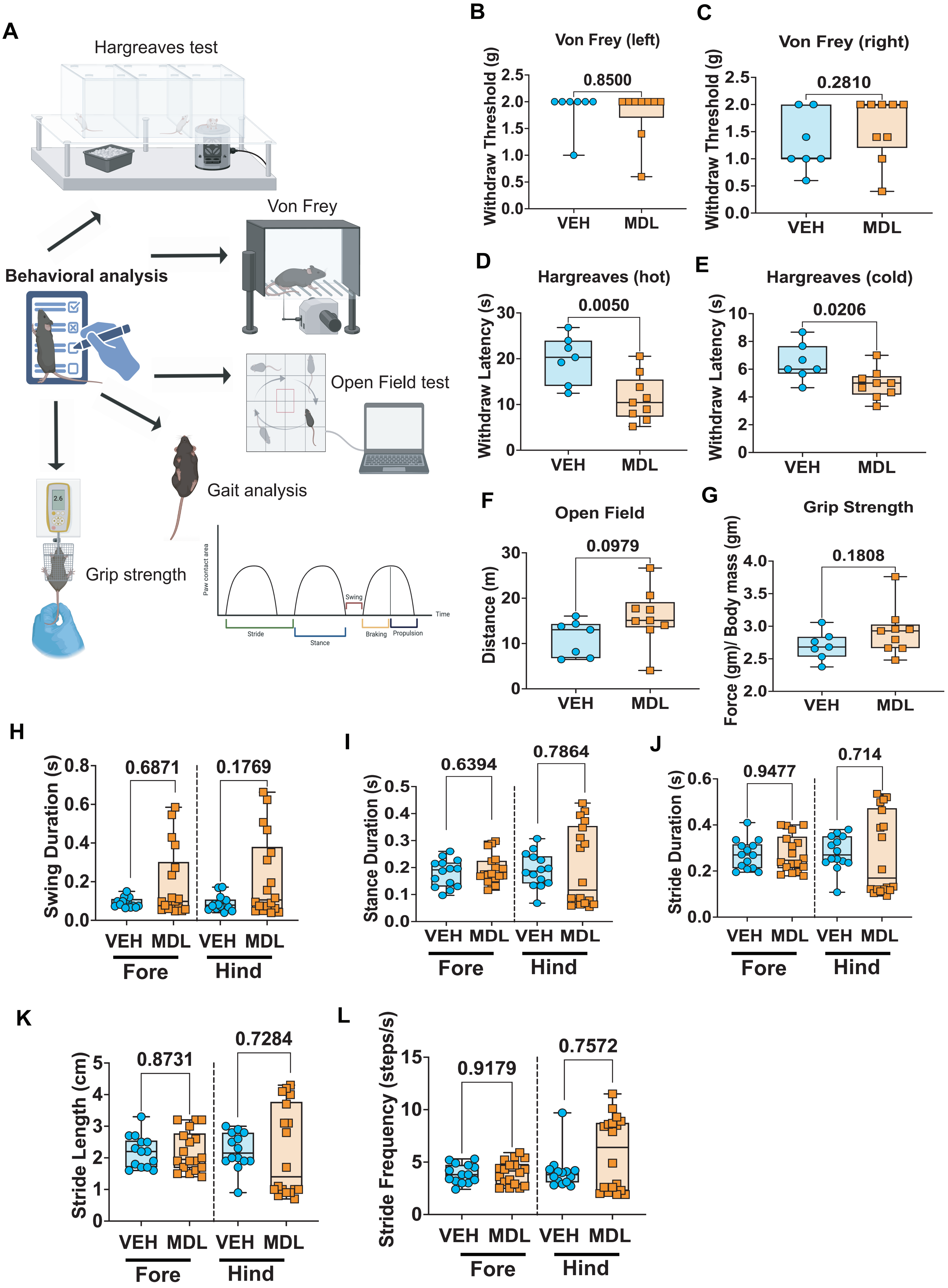

A
Hargreaves test
Von Frey
Behavioral analysis
Open Field test
Gait analysis
Grip strength
Paw contact area
Swing
Time
Stride
Stance
Braking
Propulsion
B
Von Frey (left)
Withdraw Threshold (g)
0.8500
VEH
MDL
C
Von Frey (right)
Withdraw Threshold (g)
0.2810
VEH
MDL
D
Hargreaves (hot)
Withdraw Latency (s)
0.0050
VEH
MDL
E
Hargreaves (cold)
Withdraw Latency (s)
0.0206
VEH
MDL
F
Open Field
Distance (m)
0.0979
VEH
MDL
G
Grip Strength
Force (gm)/ Body mass (gm)
0.1808
VEH
MDL
H
Swing Duration (s)
0.6871
0.1769
VEH MDL VEH MDL
Fore
Hind
I
Stance Duration (s)
0.6394
0.7864
VEH MDL VEH MDL
Fore
Hind
J
Stride Duration (s)
0.9477
0.714
VEH MDL VEH MDL
Fore
Hind
K
Stride Length (cm)
0.8731
0.7284
VEH MDL VEH MDL
Fore
Hind
L
Stride Frequency (steps/s)
0.9179
0.7572
VEH MDL VEH MDL
Fore
Hind
