## Supplemental material for "SIRT6 Activation Improves Intervertebral Disc Health in the Aging Spine"

**A** Original image for 6B

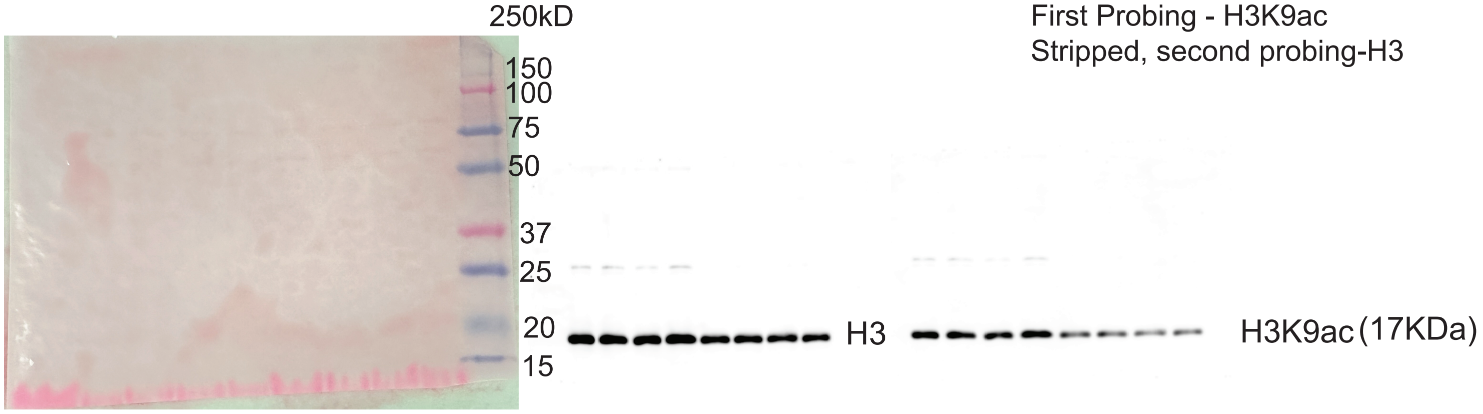


First Probing - H3K9ac
Stripped, second probing-H3

**B** Original image for 6C

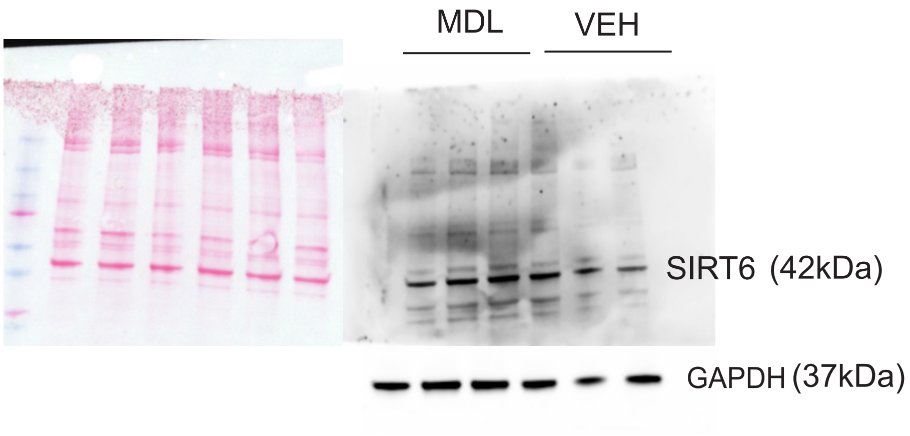


First Probing - SIRT6
Stripped, second probing-GAPDH

**C** Original image for 7A

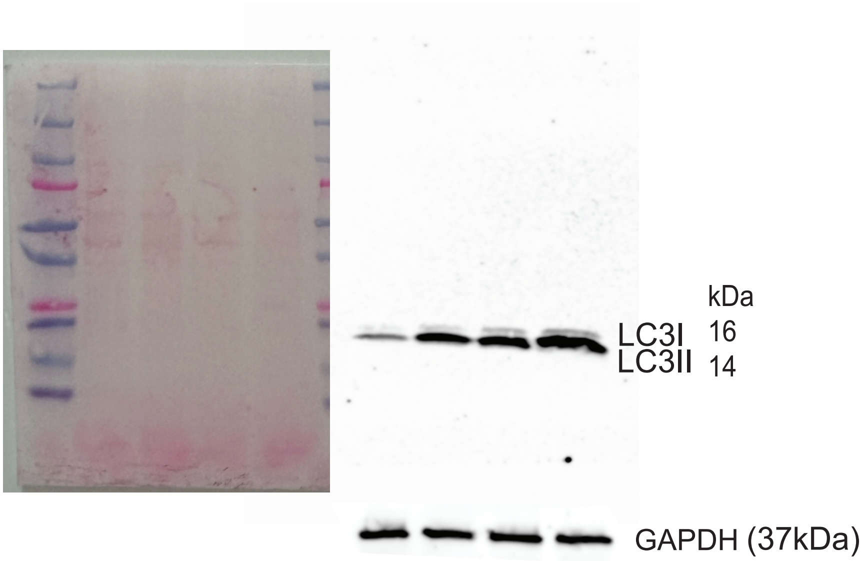


First Probing - LC3
Stripped, second probing-GAPDH
